# Massively parallel discovery of human-specific substitutions that alter neurodevelopmental enhancer activity

**DOI:** 10.1101/865519

**Authors:** Severin Uebbing, Jake Gockley, Steven K. Reilly, Acadia A. Kocher, Evan Geller, Neeru Gandotra, Curt Scharfe, Justin Cotney, James P. Noonan

## Abstract

Genetic changes that altered the function of gene regulatory elements have been implicated in the evolution of the human brain. However, identifying the particular changes that modified regulatory activity during neurodevelopment remains challenging. Here we used massively parallel enhancer assays in human neural stem cells to measure the impact of 32,776 human-specific substitutions on enhancer activity in 1,363 Human Accelerated Regions (HARs) and 3,027 Human Gain Enhancers (HGEs), which include enhancers with novel activities in humans. We found that 31.9% of active HARs and 36.4% of active HGEs exhibited differential activity between human and chimpanzee. This enabled us to isolate the effects of 401 human-specific substitutions from other types of genetic variation in HARs and HGEs. Substitutions acted in both an additive and non-additive manner to alter enhancer activity. Human-specific substitutions altered predicted binding sites for a specific set of human transcription factors (TFs) that were a subset of TF binding sites associated with enhancer activity in our assay. Substitutions within HARs, which are overall highly constrained compared to HGEs, showed smaller effects on enhancer activity, suggesting that the impact of human-specific substitutions may be buffered in enhancers with constrained ancestral functions. Our findings yield insight into the mechanisms by which human-specific genetic changes impact enhancer function and provide a rich set of candidates for experimental studies of regulatory evolution in humans.

## Introduction

Changes in developmental gene regulation are hypothesized to contribute to the evolution of novel human phenotypes such as the expansion of the cerebral cortex (Geschwind and Rakic, 2013; King and Wilson, 1975; Reilly and Noonan, 2016). However, identifying specific genetic variants that altered regulatory activity in human development remains challenging. Previous studies using comparative genomics strategies have identified two classes of regulatory elements that may encode uniquely human functions. The first class are Human Accelerated Regions (HARs), which are highly conserved across species but exhibit a significant excess of fixed sequence changes in humans (Capra et al., 2013; Lindblad-Toh et al., 2011; Pollard et al., 2006a, 2006b; Prabhakar et al., 2006). HARs are enriched near genes implicated in brain development, and several HARs have been shown to encode transcriptional enhancers with human-specific changes in activity (Capra et al., 2013; Haygood et al., 2010; Pollard et al., 2006a, 2006b; Prabhakar et al., 2006, 2008). The second class of elements are Human Gain Enhancers (HGEs), which show increased epigenetic signatures of enhancer activity in developing human tissues compared with rhesus macaque and mouse (Cotney et al., 2013; Reilly et al., 2015). Thousands of HGEs have been identified in the human embryonic cortex. Chromosome conformation studies demonstrate that HGEs and HARs target genes involved in neurogenesis, axon guidance and synaptic transmission in the developing human cortex (Rajarajan et al., 2018; de la Torre-Ubieta et al., 2018; Won et al., 2016, 2019).

The exact genetic changes that drive novel human regulatory functions in HARs and HGEs, and the effect they have on enhancer activity both individually and in combination, remain to be fully determined. One approach used to identify and characterize functional single-base variants is the Massively Parallel Reporter Assay (MPRA; Melnikov et al., 2012), in which a synthesized library of candidate regulatory elements is cloned in front of a reporter gene containing a random oligonucleotide barcode. High-throughput sequencing of the transcribed barcode collection is then used to quantify regulatory activity. A single MPRA experiment interrogates thousands of sequence changes at once, which provides the means to comprehensively analyze sequence variation within regulatory elements in a combinatorial manner. This technique has been used to quantify the effect of saturating mutations in single enhancers or to interrogate the functional effects of segregating human variants (van Arensbergen et al., 2019; Melnikov et al., 2012; Ulirsch et al., 2016).

A recent MPRA queried 714 HARs in human and chimpanzee neural progenitor cells (Ryu et al., 2018). Forty-three percent of these HARs were found to act as enhancers in this assay, two thirds of which showed differential activity between the human and chimpanzee versions. The study also dissected the sequence variation in seven of these HARs, showing that a combination of buffering and amplifying effects of single base substitutions generated a conserved or modified regulatory output (Ryu et al., 2018). This is consistent with an earlier study that partially dissected the relative contribution of single substitutions in the HAR *HACNS1*, which suggested that at least two substitutions are responsible for the human enhancer’s increased activity (Prabhakar et al., 2008). However, HARs only capture a small fraction of human-specific substitutions in the genome that may alter regulatory function. The impact of human-specific substitutions on enhancer activity overall has not been investigated.

To gain insight into this question, we used MPRAs to identify the additive and non-additive effects of over 32,000 human-specific substitutions on enhancer activity in H9-derived human Neural Stem Cells (hNSCs), a cell model system for early neurogenesis. These hNSCs model neurodevelopmental precursor cells, a stem cell niche that gives rise to cortical neurons (Gage, 2000). Regulatory changes in anterior forebrain precursors may have contributed to the expansion of the cortex during human evolution (Geschwind and Rakic, 2013). The 32,000 human-specific substitutions we queried are located in HARs or HGEs, as these include enhancers with prior evidence of human-specific function. We identified 16.3% of HARs and 43.0% of HGEs as active enhancers in human neurodevelopment, of which 31.9% and 36.4%, respectively, were differentially active between human and chimpanzee.

We then designed a second MPRA to further dissect the effects of over 1,300 human-specific substitutions within differentially active enhancers to identify 401 substitutions that modify enhancer activity in human. Some of these variants act alone, while others interact with variants nearby. Pairs of interacting substitutions were often in close proximity, in line with a model where transcription factor binding is modified in concert by multiple substitutions. We identified a suite of transcription factors implicated in enhancer activity in our assay, including cell cycle regulators. Our findings reveal mechanisms of enhancer evolution in humans and provide an entry point towards a functional understanding of the genetic changes underlying the evolution of the human neocortex.

## Results

### Interrogating human-specific substitutions within HARs and HGEs for effects on enhancer activity in a cellular model of human neurodevelopment

We used a two-stage Massively Parallel Reporter Assay (MPRA) to screen the effects of 32,776 human-specific substitutions within 1,363 Human Accelerated Regions (HARs) and 3,027 Human Gain Enhancers (HGEs) on neurodevelopmental enhancer activity (Figure 1A, Table S1). We defined a set of high-confidence human-specific substitutions (termed “hSubs”) to isolate the regulatory effects of sequence changes at sites that are otherwise unchanged throughout primate evolution. To achieve this, we restricted our screen to positions in the reference human genome (GRCh37/hg19) exhibiting a derived sequence change compared to the orthologous position in chimpanzee, orangutan, rhesus macaque and marmoset, which we required to all exhibit the same, putatively ancestral character state. We then used dbSNP (build 144) to exclude human polymorphisms (see Methods).

**Figure 1.**
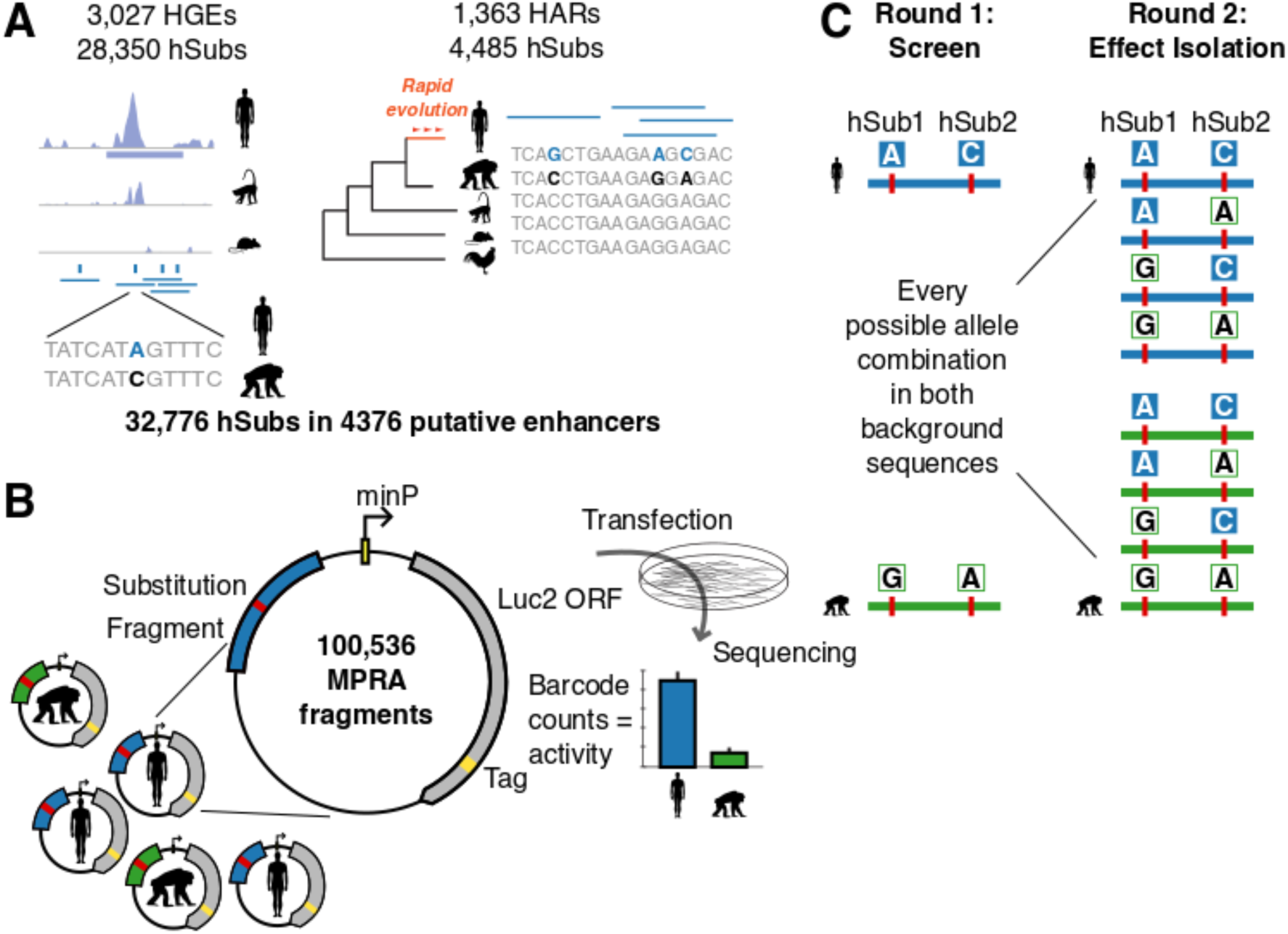
Experimental design. (**A**) We synthesized 137-bp MPRA fragments overlapping 32,776 hSubs in 3,027 HGEs and 1,363 HARs. (**B**) MPRA fragments (human in blue, chimpanzee in green) were cloned in front of a *luc2* reporter gene and a random oligonucleotide barcode tag (in yellow). Sequencing and counting barcodes provides a quantitative measure of enhancer activity. (**C**) In stage 1 of the experiment, we screened 50,268 orthologous human-chimpanzee fragment pairs. In stage 2, the impact of genetic differences within all fragments exhibiting species-specific changes in activity was dissected by testing all possible combinations of hSubs and ancestral states in both the human and chimpanzee background reference sequences.

In the first stage of our screen, we designed human–chimpanzee orthologous pairs of 137-bp MPRA fragments centered on each substitution to find active enhancer regions containing hSubs. Where two hSubs were in such close proximity that they would overlap the same fragment, we generated additional fragments centered on the midpoint between each pair of hSubs. We synthesized 100,536 fragments including pairs of human and chimpanzee orthologs (Figure 1A). To generate the MPRA library, we cloned each fragment upstream of a minimal promoter driving the expression of a *luc2* firefly luciferase ORF tagged with a random oligonucleotide barcode (Figures 1B and S1). Each MPRA fragment was linked to 80 unique barcodes on average, yielding a total library containing approximately 8 million unique molecules. We included human and chimpanzee orthologous fragments in a single library to prevent batch effects that would confound potential species differences in activity.

To carry out the MPRA, we transfected the library into hNSCs in four replicates. To quantify regulatory activity, we collected both input plasmid DNA (pDNA) and total RNA 6 hours post-transfection (Methods; Figure S8). Following cDNA synthesis, we used high-throughput sequencing to determine barcode counts in both the cDNA and pDNA fractions (Methods). Barcode counts were highly similar across replicates (Spearman’s rank correlation for barcode counts, *ρ* = 0.87–0.89 for pDNA and *ρ* = 0.81–0.85 for cDNA, *P* < 1 · 10^−300^), especially after averaging barcodes per fragment (*ρ* = 0.87–0.89 for pDNA and *ρ* = 0.84–0.87 for cDNA, *P* < 1 · 10^−300^). We found that fragments with small barcode numbers showed high variance in barcode representation across replicates. However, barcode representation stabilized at 12 or more barcode tags per fragment, so we excluded all fragments with pDNA barcode counts below this threshold (Figure S2). 78.1% of fragments passed this threshold (78,487 of 100,536 single fragments; hereafter called ‘measured fragments’). Measured fragments had an average of 69.3 barcodes (SD = 62.4; 95% range, 13–233).

Barcode counts were highly correlated overall between pDNA and cDNA fractions in each replicate (*ρ* = 0.81–0.84 for barcode replicates and *ρ* = 0.79–0.81 for fragment replicates, *P* < 1 · 10^−300^). However, we identified a population of fragments from each species that exhibited increased levels of cDNA-derived barcode counts, suggesting they act as transcriptional enhancers (Figures 2A–B). For each barcode, we calculated the ratio of cDNA over pDNA counts, which we refer to as ‘activity’. We then quantified fragment activity as the log_2_ mean of activities of all barcodes assigned to that fragment. We found fragment activity to be robust between replicates (*ρ* = 0.73–0.78, *P* < 1 · 10^−300^). To identify fragments displaying significant enhancer activity, we used one-tailed *t* tests to compare each fragment’s activity against the rest of the library (Methods). In addition, we used Mann-Whitney *U* tests to account for fragments with non-normal barcode distributions. We also required that every replicate had higher cDNA than pDNA counts and that the log_2_ activity averaged over all replicates was larger than 0.1. The union of all fragments with positive test results is our initial set of active fragments.

**Figure 2.**
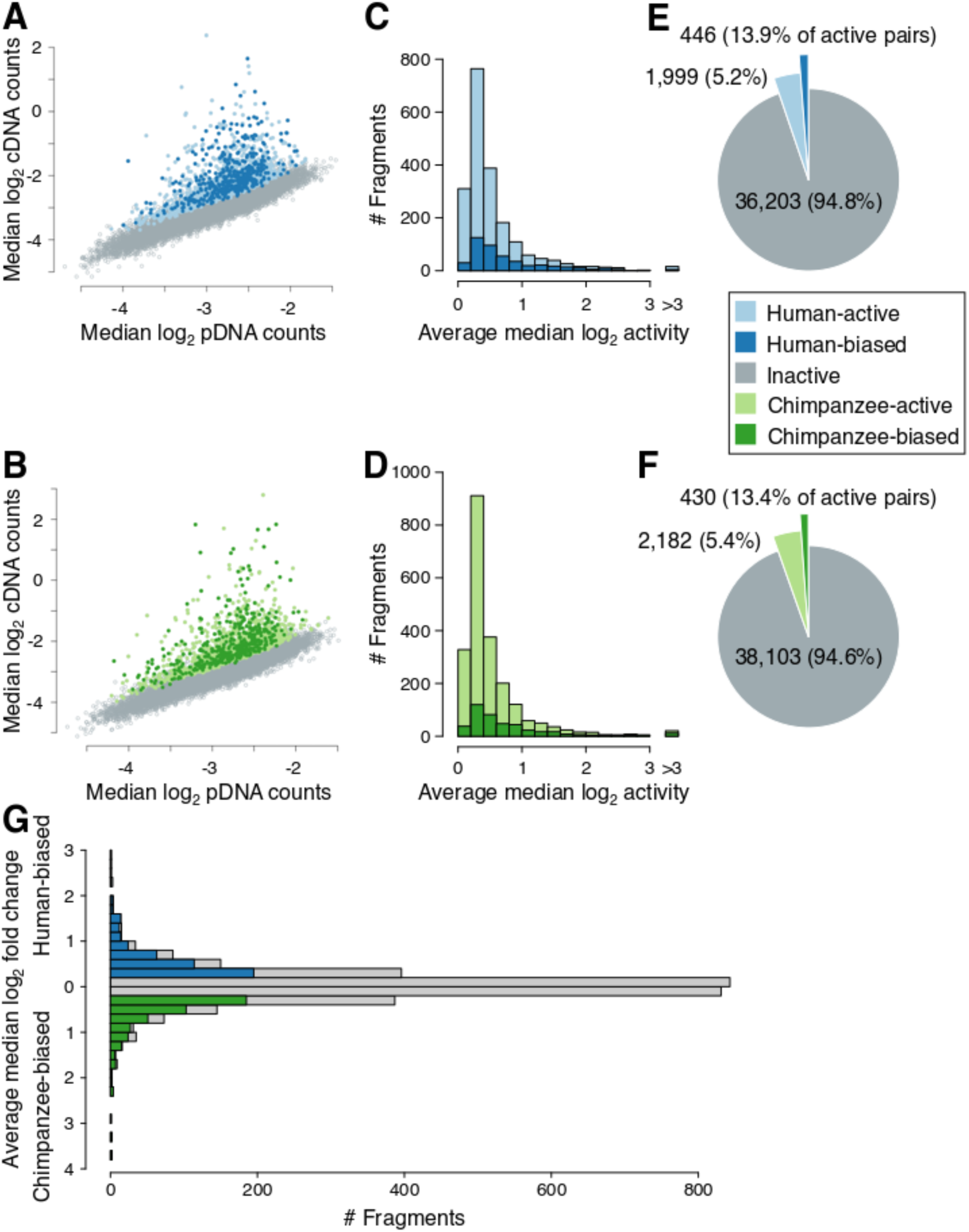
Quantifying enhancer activity using MPRA. (**A**–**B**) Comparing MPRA barcode counts in pDNA vs cDNA fractions identifies fragments showing increased numbers of cDNA counts, indicative of enhancer activity. (**A**) Active human fragments are shown as light blue points and differentially active human fragments are shown in dark blue. (**B**) Active chimpanzee fragments are shown as light green points and differentially active chimpanzee fragments are shown in dark green. (**C**–**D**) The distribution of enhancer activity in all active and differentially active fragments in human (**C**) and chimpanzee (**D**). (**E**–**F**) The number of active and differentially active fragments detected from human (**E**) and chimpanzee (**F**). (**G**) The distribution of differences in activity between all active human and chimpanzee fragments. Human and chimpanzee fragments showing significant increases in activity are labeled in dark blue or dark green, respectively. Fragments showing non-significant activity differences are shown in gray.

We identified 4,181 fragments (5.3% of all measured fragments; Table S2A) that showed increased transcriptional activity (Figures 2C–F). Most of these fragments showed modest activity: 11.7% of active fragments showed an average activity higher than 2 (meaning their cDNA production was twice as high as their pDNA input). Active fragments were more commonly found in regions with evidence of regulatory activity in hNSCs based on ATAC-seq or H3K27ac peaks (Table S3). While HGEs contained a larger total number of measured (36,656 vs. 7,884) and active fragments (2,776 vs. 457) than HARs, the relative proportion of active fragments was higher in HARs than in HGEs (17.3% vs. 13.2%). However, we may be underestimating the proportion of functional sequences in HGEs since we are focusing only on regions that include human-specific substitutions. In contrast, HARs were sampled more densely due to their deep conservation and high substitution density.

We then sought to identify orthologous human and chimpanzee fragments that showed differential enhancer activity. We applied two-tailed *t* and *U* tests to compare the activity of all 3,219 measured fragment pairs that had at least one active fragment in either ortholog (Material and Methods). We also required that fragment activity was biased in the same species direction in all replicates, and imposed a minimum average difference threshold > log_2_ 0.1. We identified 876 differentially active fragment pairs (27.2% of all active pairs; Figures 2E–F, Table S2A). Most of these showed modest changes in activity between species (mean fold change = 1.61, 95% range, 1.16–3.29; Figure 2G). However, 114 fragments (13.0%) showed a fold change larger than 2 and 15 fragments (1.7%) showed a fold change larger than 4 (Figure 2G). We found that differentially active fragments were not more likely to be human-biased than chimpanzee-biased (*P* = 0.66, two-tailed binomial test). Differentially active fragments were also more frequently found in regions with evidence of regulatory element activity based on chromatin signatures in hNSCs than inactive fragments (Table S3).

We next determined the distribution of active and differentially active fragments within HARs and HGEs. We identified a HAR or HGE as active if it contained at least one active fragment and as differentially active if it contained at least one differentially active fragment. We found that 16.3% of HARs and 43.0% of HGEs were active and that 5.2% and 15.7%, respectively, were differentially active (Figures 3A–B). This included 30 human-biased HARs, 228 human-biased HGEs, 36 chimpanzee-biased HARs and 213 chimpanzee-biased HGEs. An additional 27 HGEs were both human- and chimpanzee-biased in different parts of their sequence. A substantial fraction of the putative enhancers we studied thus showed MPRA activity even though the overall number of fragments that showed activity was small, consistent with previous MPRA studies (Tewhey et al., 2016; Ulirsch et al., 2016). As was the case at the fragment level, there was no preference for enhancer activity to be biased in the direction of either species (Figures 3C–D). Overall, our MPRA captures changes in enhancer activity in NSCs that potentially reflect species-specific regulatory differences in human corticogenesis.

**Figure 3.**
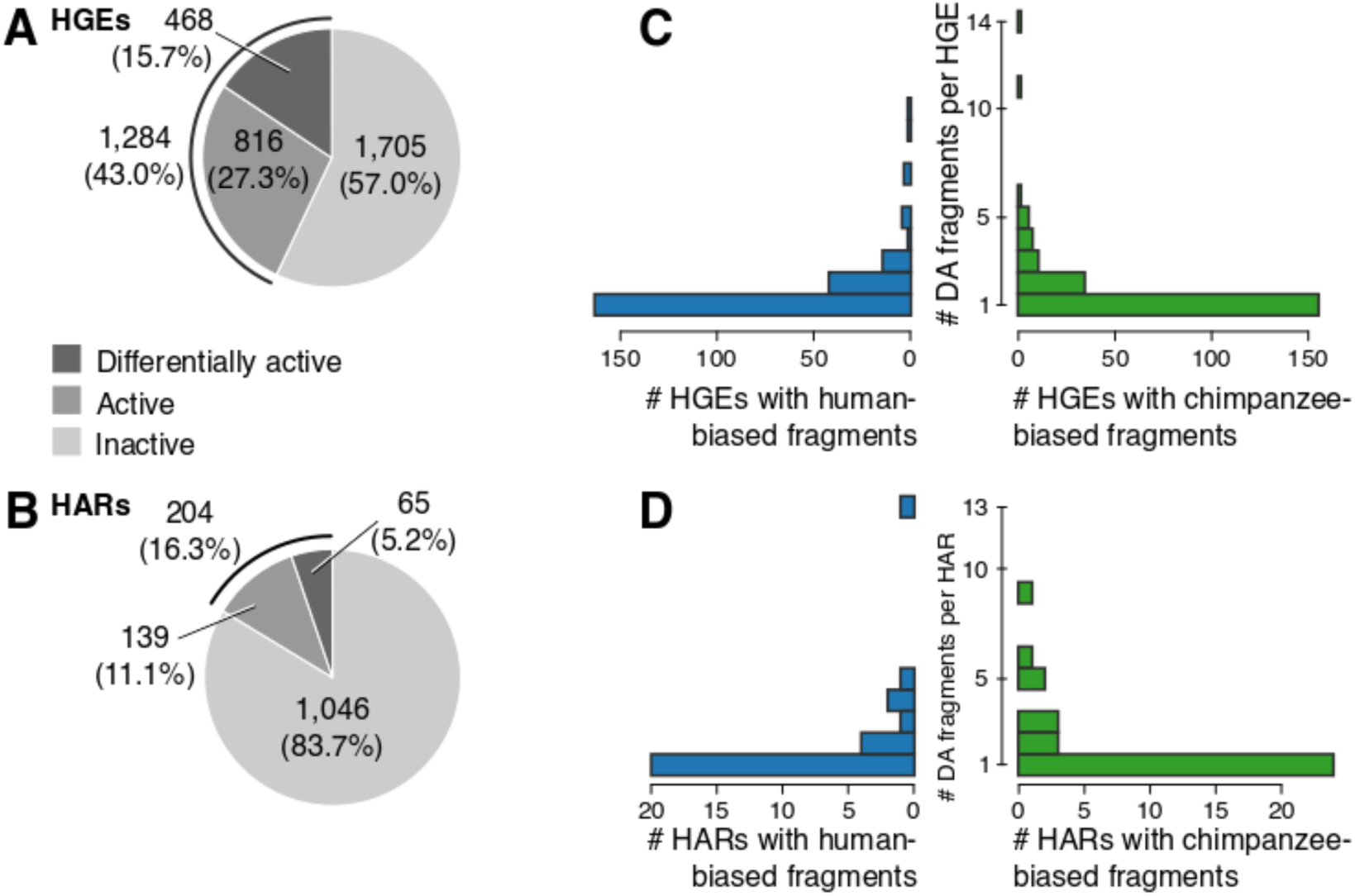
Active and differentially active HARs and HGEs identified by MPRA. (**A**–**B**) The proportion of inactive, active and differentially active HGEs (**A**) and HARs (**B**). (**C**–**D**) The distribution of the number of differentially active fragments per HGE (**C**) and HAR (**D**). Most species-biased HGEs and HARs contain only one differentially active MPRA fragment.

We compared our findings with those from a recent study (Ryu et al., 2018) that screened 714 HARs (Lindblad-Toh et al., 2011; Pollard et al., 2006a) in an MPRA using a lentiviral integration design in human and chimpanzee neural progenitor cells at two different time points after neural induction. While this study detected a larger number of active HARs, likely due to different criteria used for identifying significantly active fragments (see Discussion), there was good agreement between the findings in both studies. Of the 76 HARs active in our study and tested in both studies, 51 (67%) were found active in both. Of the 17 HARs that contained differentially active fragment pairs in our study and were tested in both studies, 14 (82%) showed differential activity in both.

### Isolating the effects of human-specific substitutions in HARs and HGEs

We designed a second MPRA experiment to isolate the specific effects of each hSub in differentially active fragments. In addition, we included validation fragments (identical copies) for all fragments that were active in the first round (including both fragments of a pair if only one was active). This second MPRA library contained a total of 24,543 fragments (of which 22,860 were measured with ≥12 barcodes) with a larger number of barcodes per fragment compared to the first MPRA (mean = 179.5 barcodes per measured fragment; 95% range = 16–808). We transfected this library into hNSCs in two replicates and measured activity by sequencing pDNA and cDNA barcode libraries.

The second round of our MPRA replicated the results from the first round well. Fragment activity was highly correlated between the two MPRA rounds (Pearson’s *ρ* range, 0.88–0.92, *P* < 1 · 10^−300^), similar to correlations of replicates within each round of MPRA (*ρ*, 0.91 [round 1], 0.94 [round 2], *P* < 1 · 10^−300^). In the second round, we included a set of negative control fragments with comparable sequence content as experimental fragments from genomic regions that exhibit no evidence of enhancer activity (Methods). We then tested the activity of experimental fragments against these negative controls. We chose this approach because the second library consisted solely of fragments with prior evidence of activity. This was in contrast to the first MPRA round, in which most fragments were expected to be inactive and could thus be used as a null distribution for identifying significantly active fragments. Using these negative controls as a background set, we used one-tailed *t* tests to identify fragments with significant activity. Using stricter criteria for identifying significantly active fragments in this experiment than in the first MPRA (see Methods), we found that 66.3% of the fragments that were active in round one and measured in both experiments replicated their activity as enhancers in the second MPRA (Table S2B). Of the fragment pairs that were significantly differentially active in round one and measured again in round two, 89.1% were active, of which 54.7% were differentially active (two-tailed *t* test; Table S2B).

In the first round of MPRA, we identified fragments that were differentially active between species. These fragments may contain both hSubs and additional background genetic variation, including human polymorphisms and substitutions that did not meet our strict criteria, that is of less interest for understanding human regulatory evolution. To isolate the specific effects of individual hSubs from this background variation, we generated fragments containing all possible combinations of the hSub and the corresponding chimpanzee allele on both the human and the chimpanzee background sequence for all differentially active fragment pairs (Figure 1B). We interrogated 1366 hSubs using 14,429 combinatorial fragments. We then employed an ANOVA modeling scheme to identify hSub-specific effects on enhancer activity. This model distinguished i) additive effects of hSubs; ii) additive background variation effects; iii) interactions between pairs of hSubs; and iv) interactions between hSubs and the background. We identified 401 hSubs that showed significant effects on fragment activity, either alone (additive; n = 315) or in combination with other hSubs or background variation (interactive; n = 120; note that a subset of individual hSubs can be scored as additive or interactive in two different fragments; Figure 3). We calculated hSub effect sizes as the fold change normalized by the standard deviation (SD).

There was no significant species bias in the direction or size of hSub effects (Mann-Whitney *U* = 13,137, *P* = 0.35 for additive effects; *U* = 1705, *P* = 0.81 for interactive effects; Figure 4E). For two-way interactive effects, we extracted the larger of the two effect sizes, i.e., the larger effect size compared to the two reference states (hSub or background variation). This state is where interactive effects exert their main function, and most of the remaining effect sizes were negligibly small (65% of the smaller effect sizes were < 0.2 SDs). Interactive effect sizes were significantly larger than additive effect sizes (additive: mean = 0.43 SDs; interactive: mean = 0.65 SDs; *U* = 13,666, *P* = 8.0 · 10^−6^). However, in both cases effect sizes were generally small (quartile range for additive effects, 0.16–0.55; for interactive effects, 0.26–0.75). Despite this, we did identify hSubs with large effects. Of the 315 hSubs with additive effects, 24 were > 1 SD and two were > 2 SDs (Table S4). Of the 120 hSubs with two-way interactive effects, 16 effects were > 1 SD and five were > 2 SDs. The maximal effect sizes observed were associated with interactive effects; the most human-biased effect was 3.36 SDs and the most chimpanzee-biased effect 4.01 SDs. The largest additive effects were smaller at 2.26 (human-biased) and 2.90 (chimpanzee-biased) SDs.

**Figure 4.**
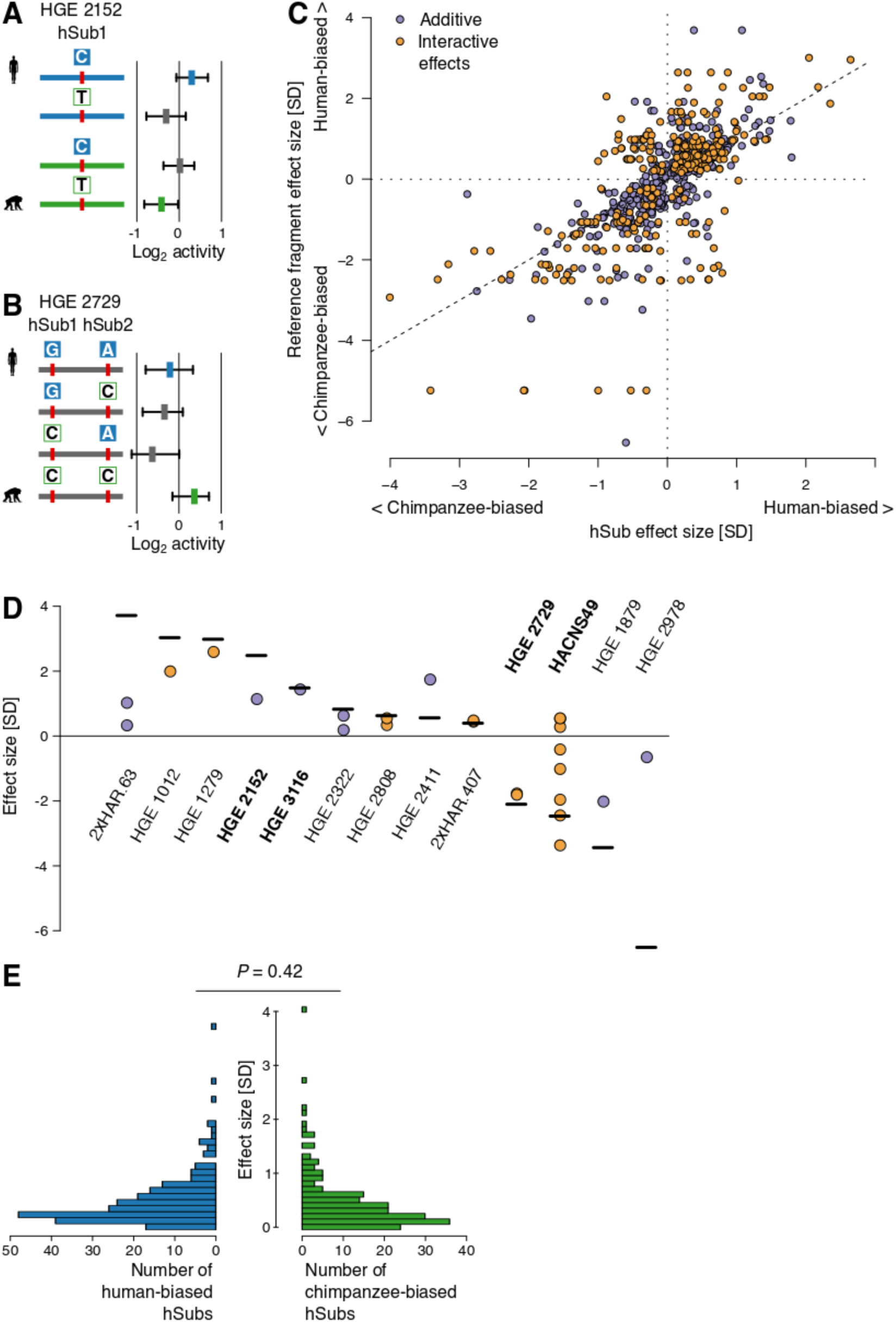
Dissecting additive and interactive effects of human-specific substitutions on enhancer activity. (**A**) The single hSub in this locus shows an additive human-bias, enhanced by the background variation’s human-biased effect. The boxplots at the right show the median and quartiles of barcode distributions for each fragment. The blue box shows the activity of the human reference fragment, the green box the activity of the chimpanzee reference fragment and gray boxes the activities of synthetic intermediates. Human hSub allele states are shown as filled blue boxes, and chimpanzee allele states are shown in green. The bar corresponding to each fragment is colored based on whether the human (blue) or chimpanzee (green) reference sequence includes additional background variation. Grey indicates that the human and chimpanzee references are identical other than the hSubs analyzed. (**B**) An example of an interaction effect between two hSubs. The alleles containing either both chimpanzee hSub states (CC) or both human hSub states (GA) show higher enhancer activity than either intermediate. However, the human allele is less active than the chimpanzee allele, suggesting loss of function. Colors correspond to those in **A**. See Figure S3 for additional examples. (**C**) All additive and interactive hSub effect sizes plotted against the reference allele effect size. Each hSub is indicated by a colored circle. Positive values on either scale indicate human-biased activity, while negative values indicate chimpanzee-biased activity. All hSubs from the same reference fragment will have the same value on the Y-axis. Points along the diagonal indicate hSub effects that align with the reference fragment effect. (**D**) Comparisons of reference fragment effect sizes with the hSubs within them. Out of the 404 fragments with significant hSub effects, the top and bottom three and additional example loci are shown. Each locus is labeled by its HAR or HGE designation. The examples mentioned in the main text and those shown in panels **A** and **B** are labeled in bold text. The total reference fragment effect size is shown by a horizontal line. In cases where hSub effects do not add up to the fragment effect size, background variation and statistical noise make up the remainder of the effect. Figure S4 shows all fragments. (**E**) Effect size distributions of human and chimpanzee hSubs. Each horizontal bar shows the number of hSubs with a given effect size, plotted on the Y-axis. A Mann-Whitney *U* test indicates no difference in distribution.

An example of an hSub with an additive effect on enhancer activity is shown in Figure 4A. This fragment contains one hSub and additional background sequence differences between the human and chimpanzee alleles. The T->C hSub has a major effect on fragment activity independent of the background sequence differences: the human-specific C allele is more active than the ancestral T allele. The background variation also contributes to differences in overall fragment activity, with the human allele being more active than the chimpanzee allele. These results illustrate the value of isolating the effects of *bona fide* human-specific substitutions, which are most relevant for understanding and characterizing regulatory changes that underlie uniquely human biology.

An example of hSubs interacting to alter enhancer activity is shown in Figure 4B. This fragment contains two hSubs and no background variation. The human (GA) and chimpanzee (CC) reference alleles are both more active than either of the synthetic intermediates (GC or CA). This shows how the activating effect of one hSub allele may depend on the allele state of another hSub. Only a specific combination of allele states leads to high activity, while other combinations lead to a reduced activity. The chimpanzee allele is the most active allele overall. Three more complex examples are shown in Figure S3.

Taken together, isolating the effects of single hSubs reveals how multiple hSubs may combine to alter enhancer function (Figures 4C–D and S4). hSub effects may largely align with the resulting fragment effect. In the simplest case, a single hSub in the absence of any background variation will cause the entire change in fragment activity (e.g., HGE 3116 in Figure 4D). In more complex cases, individual hSub effects may point in the opposite direction of the difference in fragment activity, thereby buffering the overall fragment effect (e.g., *HACNS49* in Figure 4D). Such effects are then compensated for by other hSubs or by the background variation in the same fragment. Fragments that contained larger numbers of hSubs often showed more complex interactions between hSubs and the background, as can be seen by the large spread of hSub effects within some fragments (e.g., *HACNS49* in Figure 4D). Fragments containing more hSubs tended to have a greater overall effect size (Spearman’s rank correlation *ρ* = 0.12, *P* = 0.0080).

### Effect sizes of human-specific substitutions differ between HGEs and HARs

We found that hSubs in HGEs exhibited significantly larger overall effect sizes than hSubs in HARs (HGEs: mean = 0.51 SDs; HARs mean = 0.38 SDs; Mann-Whitney *U* = 19,211; *P* = 1.1 · 10^−7^; Figure 5B). We considered several possible mechanisms that could account for this finding. First, HARs are more often inactive than HGEs in our MPRA. HGEs were defined based on epigenetic signatures of enhancer activity in the developing human cortex at time points when substantial numbers of neural stem cells are present (Reilly et al., 2015). In contrast, HARs were defined based on a significant excess of human-specific substitutions in otherwise deeply conserved regions, without reference to any potential biological function (Lindblad-Toh et al., 2011; Pollard et al., 2006a, 2006b; Prabhakar et al., 2006). To investigate this, we categorized HARs according to whether or not they reside in H3K27ac ChIP-seq peaks in hNSCs (Methods). We did not find a significant difference between hSubs in active and inactive HARs, although the sample size was small (*U* = 817; *P* = 0.57; Figure 5B).

**Figure 5.**
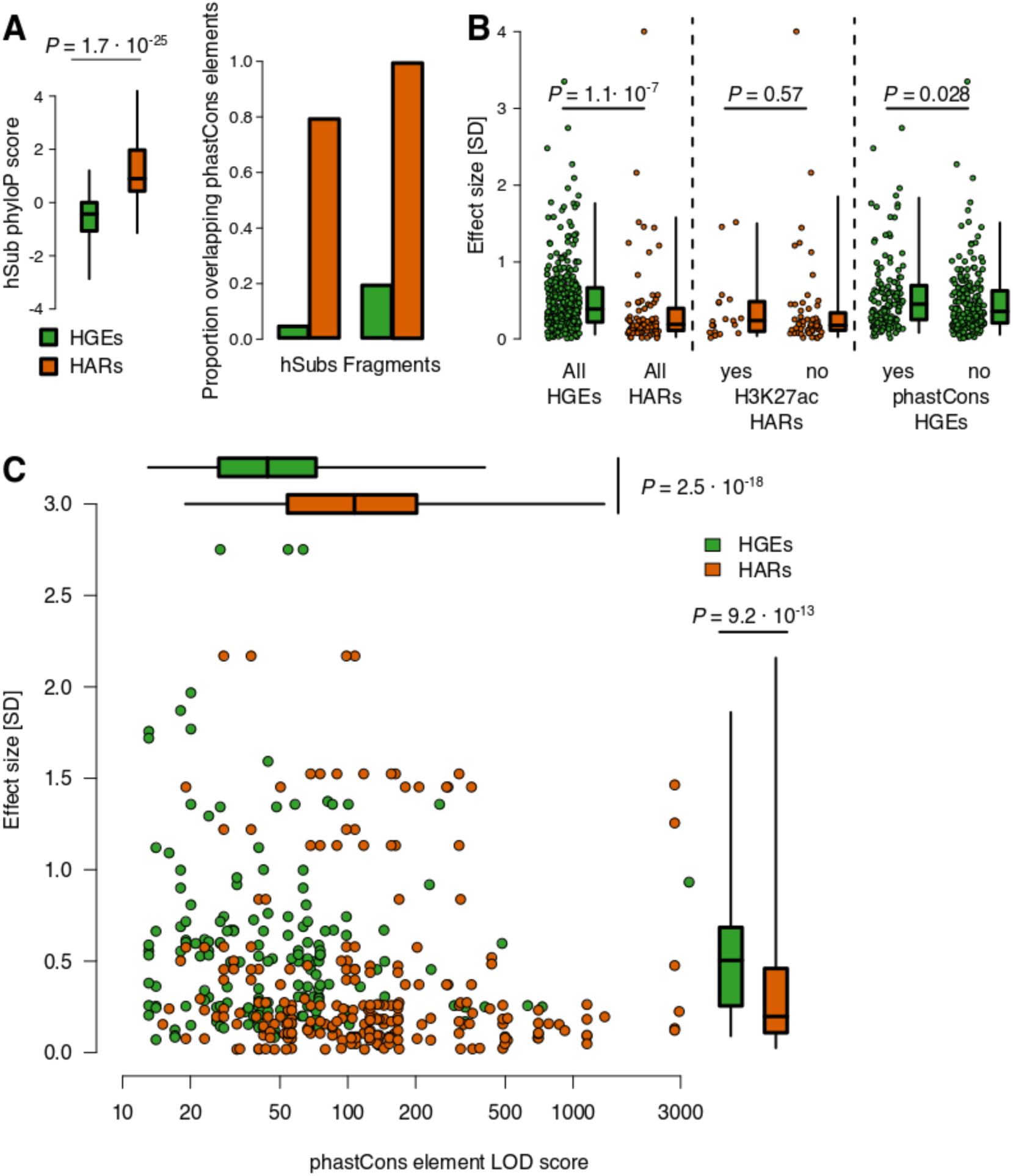
Substitutions in HARs and HGEs differ in their effects on enhancer activity. (**A**) Both hSubs and fragments in HGEs (green) and HARs (orange) differ in their level of constraint. (**B**) hSubs in HGEs have significantly larger effect sizes than hSubs in HARs (left panel). However, hSubs in HARs that show evidence of activity in hNSCs based on chromatin signatures do not have significantly larger effects than hSubs in inactive HARs (middle panel). hSubs in fragments with evidence of constraint in HGEs show larger effects than hSubs in unconstrained fragments (right panel). (**C**) HARs (orange) and HGEs (green) differ in effect size of their hSubs (shown on the Y-axis in the scatterplot and in the box plots on the right) and in evolutionary conservation (measured as the LOD score of phastCons elements overlapping MPRA fragments; shown on the X-axis in the scatterplot and in the box plots at the top of the figure), but these two aspects are uncorrelated.

Second, HARs are substantially more conserved than HGEs, and are likely to encode regulatory functions that are still under some degree of constraint in humans. The effect of hSubs in HARs may be buffered due to this prior constraint. However, HGEs include both constrained and unconstrained sequences, and hSubs in unconstrained regions may introduce novel enhancer activity without disrupting ancestral functions. To evaluate this hypothesis, we compared the distribution of constraint in HARs and HGEs using phastCons and phyloP (Pollard et al., 2010; Siepel et al., 2005). We observed a higher phyloP constraint score for hSubs in HARs than in HGEs (mean phyloP_HGE_ = −0.57 vs mean phyloP_HAR_ = 1.20; *U* = 2,490; *P* = 6.2 · 10^−25^; Figure 5A). Similarly, hSubs and fragments in HARs overlapped a higher proportion of constrained elements detected by phastCons than in HGEs (5.5% [hSubs] and 20.2% [fragments] for HGEs, 79.8% and 100% for HARs; Figure 5A).

From the observation that hSubs in HARs showed smaller effect sizes than those in HGEs, we predicted that hSubs in constrained HGEs would show smaller effect sizes than those in unconstrained HGEs. However, we found the opposite to be true: hSubs in fragments that overlap phastCons elements showed a significantly larger effect size than those in fragments that did not overlap a phastCons element (*U* = 17,625, *P* = 0.012; Figure 5B). While we detected an effect size difference between constrained and unconstrained HGEs, we did not find a correlation between hSub effect size and strength of constraint (Figure 5C). In summary, while sequence constraint seems to play a role in determining hSub effect size, it affects hSub effect size differently in HGEs and HARs.

### Human-specific substitutions alter predicted transcription factor binding in HARs and HGEs

To identify transcription factors (TFs) potentially driving human-specific enhancer activity in in our MPRA, we mapped all vertebrate transcription factor binding site (TFBS) motifs from the JASPAR core database onto the human and chimpanzee genomes (see Methods). We first tested for enrichment of TFBS predictions within orthologous sequences of entire MPRA fragments, including both hSubs and background variation, to identify TFBS motifs that were enriched among active or differentially active fragments. We identified 65 TFBS motifs that were enriched among all active fragments relative to all measured fragments (resampling test, *P*_BH_ < 0.05; Table S5). With the notable exception of the cell cycle control factor TP53, we found no further enrichment among TFBSs in differentially active, human- or chimpanzee-biased fragments (Table S5, Figure S5). This suggests that there is a subset of transcription factors implicated in driving enhancer activity in our assay, and that there is no more specific subset driving species-biased activity.

We next investigated if hSubs specifically altered TFBSs between human and chimpanzee enhancer sequences. The number of TFBSs overlapping hSubs within differentially active fragments was not sufficient for an informative enrichment test. However, since overall TFBS content did not differ significantly between active and differentially active fragments, comparing differences in TFBS representation between the human and chimpanzee sequences of active fragments may identify changes in TFBS recruitment that alter enhancer activity. To evaluate this, we extracted TFBS predictions overlapping hSubs in active MPRA fragments and tested for overrepresentation of TFBS motifs predicted in the human or chimpanzee orthologs relative to the union of both sets. We found 11 TFBSs enriched among human sequences and 15 enriched among chimpanzee sequences (Table S5). Nine of the TFBSs enriched in human sequences are basic helix-loop-helix transcription factors (both Hairy-related and bHLH-ZIP factors), and five of them were also among those enriched in active fragments. Taken together, these analyses provide the basis to identify individual hSubs that putatively change TF binding in HARs and HGEs.

Three quarters of the 401 hSubs with significant individual regulatory effects overlapped predicted TFBSs, including many of those enriched in active fragments. Of the hSubs that showed the largest effect sizes, one example is shown in Figure 6A. In this case, an hSub (chr5:108,791,729) generates a predicted TFBS for AP-1 TFs (FOS and JUN proteins) in the human sequence. Neither of those TFs has a predicted TFBS at the orthologous chimpanzee site. The hSub increases fragment activity by a factor of 1.98 (human-biased effect size = 1.8 SDs) in the human compared to the chimpanzee ortholog (Figure 6B). AP-1 typically binds to promoter and enhancer sequences to activate target gene expression (Chiu et al., 1988), providing a potential mechanism to explain the increased activity of this enhancer in our MPRA. AP-1 has also been suggested to control important cell cycle regulators such as Cyclin D1 and TP53 (Shaulian and Karin, 2002).

**Figure 6.**
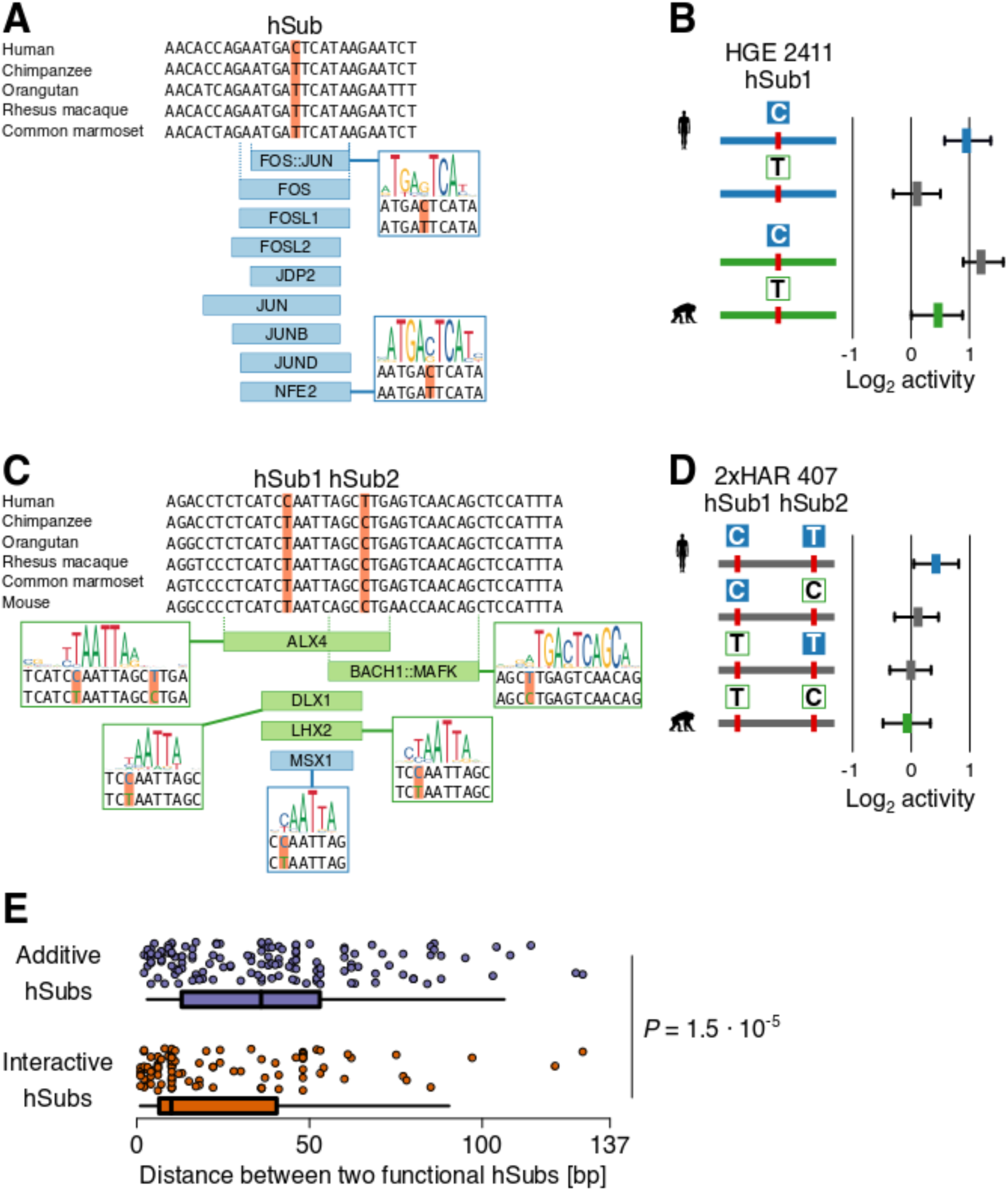
Changes in predicted transcription factor binding sites due to hSubs that alter enhancer activity. (**A**) Five-primate alignment over an hSub in HGE 2411. The alignment was derived from the UCSC 100-way Multiz alignment in GRCh37/hg19. Nine TFs are predicted to bind the human, but not the chimpanzee ortholog (TFBSs predicted to show increases in affinity are highlighted in blue). (**B**) MPRA activity of the hSub shown in **A**, as well as the activities of the chimpanzee ortholog and synthetic intermediates. The figure is labeled as in Figure 4A. (**C**) Alignment of a pair of hSubs in 2xHAR 407 that overlaps a cluster of TFBSs (blue, stronger predicted binding in human; green, stronger predicted binding in chimpanzee). See Figure S6 for additional TFBSs in this locus that are not affected by the hSubs. (**D**) MPRA activity of the two hSubs shown in **C** and the corresponding chimpanzee ortholog and synthetic intermediates. (**E**) Distribution of physical distances between pairs of additive (top, in blue) and interactive (bottom, in orange) hSub with regulatory effects. BH-corrected P-values were calculated using a Mann-Whitney *U* test.

We then examined fragments with multiple hSubs that altered predicted TFBS content. In the example shown in Figure 6C, two closely spaced hSubs overlap 14 predicted TFBSs and are in close proximity to an additional three. Four of the TFBSs (*ALX4*, *BACH1*::*MAFK*, *DLX1* and *LHX2*) show increased predicted binding affinity for the chimpanzee sequence, one shows increased predicted affinity for *MSX1* in human (Figure 6C), while the rest show only marginal change in their predicted affinity for either allele (Figure S6). Both hSubs by themselves increase MPRA activity in human over chimpanzee (human-biased effect size of hSub1 = 0.22 SDs and of hSub2= 0.20 SDs). Their combined effect on MPRA activity, however, is stronger than what would be expected from their individual effects, meaning their small individual effects combine to produce a stronger interactive effect (effect size = 0.58 SDs). BACH1::MAFK and DLX1 are known transcriptional repressors, suggesting the hSubs in this example may generate increased activity in human by disrupting recruitment of these factors (Chiba et al., 2003; Kitamuro et al., 2003; Sun et al., 2002).

These findings suggest that the interactive effects of hSubs on enhancer activity could be due to multiple hSubs altering the same or physically adjacent TFBSs. Supporting this, interacting hSubs were on average more closely spaced than additive hSubs (Mann-Whitney *U* = 9362, *P* = 1.5 · 10^−5^; Figure 6E). We found that 51.5% of interacting hSubs were clustered within 10 bp of each other, while additive hSubs did not show such clustering (19.7% of additive hSubs were within 10 bp). While we also found some distant interacting hSubs, the fact that many interacting hSubs were relatively close to their interaction partners suggests a mechanism in which two hSubs together influence transcription factor binding.

### Repressing enhancers showing human-specific activity perturbs expression of their target genes

To identify genes potentially regulated by enhancers with human-specific gains in activity, we used a high-resolution chromatin contact map generated in induced pluripotent stem cell-derived neural progenitor cells (Rajarajan et al., 2018). This dataset allowed us to infer regulatory interactions between differentially active enhancers and their putative target genes. We identified contacts between 32 differentially active enhancers including 21 hSubs with significant effects on activity and 54 genes (Tables S6 and S7). These genes were enriched for Gene Ontology categories related to cell cycle, proliferation, development and gene regulation (Table S8).

We then used CRISPRi in hNSCs to repress three differentially active enhancers and measured the resulting impact on expression of their predicted target genes using qRT-PCR. In two cases, *SDC2* and *AGAP1*, repression of the candidate enhancer significantly altered gene expression in one of two biological replicates, providing limited evidence that the targeted enhancer regulates each gene (Figure 7). We observed a stronger and significant effect of enhancer repression on the expression of *DPYSL5* in two biological replicates, suggesting the enhancer we targeted substantially contributes to *DPYSL5* regulation in hNSCs.

**Figure 7.**
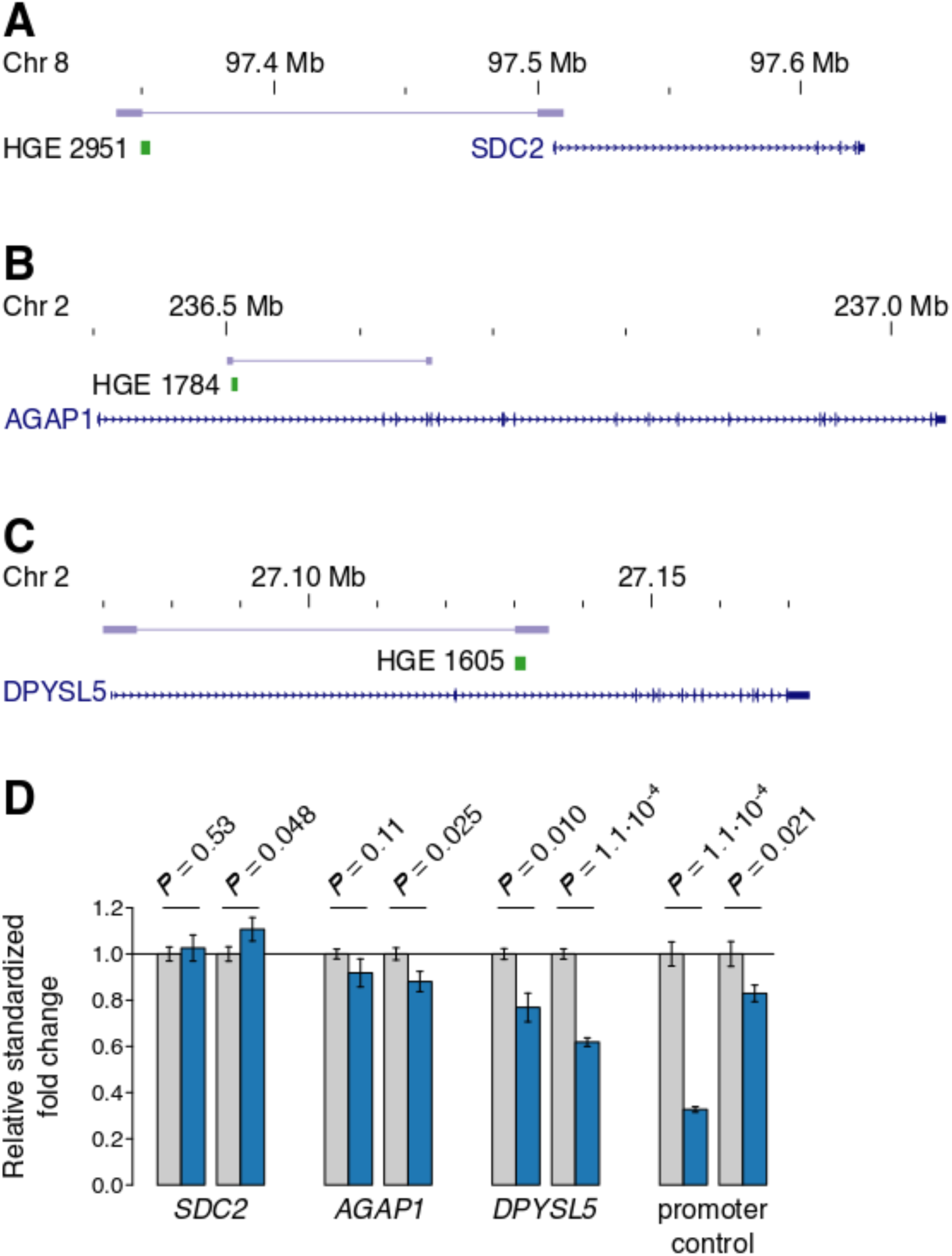
Repression of species-biased enhancers affects target gene expression. (**A-C**) Three gene-enhancer interactions targeted by CRISPRi in human neural stem cells. Each targeted enhancer is labeled and shown in green. Each gene model is shown in dark blue. The long-range contacts identified by (Rajarajan et al., 2018) are shown in light blue, with the inferred contact points shown as light blue rectangles. (**A**) HGE 2951 contacts the promoter of *SDC2*, 150 kb upstream; (**B**) HGE 1784 resides in an intron of *AGAP1*; (**C**) HGE 1605 contacts the promoter of *DPYSL5*, located 60 kb away. (**D**) qRT-PCR measurements of gene expression in hNSCs targeted by CRISPRi-sgRNAs (blue) compared to a scrambled guide control (gray). Gene expression of target genes is affected by inactivating the enhancer using CRISPRi in one (*SDC2*, *AGAP1*) or two (*DPYSL5*) CRISPRi replicates. We also show a positive control targeting the promoter of *IRX2*. BH-corrected *P* values were obtained using two-tailed *t* tests.

## Discussion

Identifying genetic changes that altered molecular functions in human evolution is the essential first step toward understanding the origins of uniquely human traits. Here we used MPRAs to screen over 32,000 human-specific substitutions (hSubs) for their effects on the activity of putative transcriptional enhancers implicated the evolution of the human cortex. We assayed 4,376 Human Accelerated Regions (HARs) and Human Gain Enhancers (HGEs) and identified members of each class that act as enhancers in our assay, as well as enhancers with differential activity between the human and chimpanzee orthologs.

HGEs were more frequently active than HARs. This is consistent with the fact that HGEs were identified based on epigenetic signatures of enhancer activity in the human, rhesus macaque and mouse developing cortex, while HARs were originally identified using a comparative genomics approach without prior evidence of function (Lindblad-Toh et al., 2011; Pollard et al., 2006a, 2006b; Prabhakar et al., 2006; Reilly et al., 2015). However, the proportion of differentially active enhancers was similar in each class, indicating that HARs and HGEs both include enhancers with novel activity in humans.

We then isolated the effects of 1,366 hSubs in differentially active fragments from each other and from background variation. This identified 401 hSubs with significant individual effects on enhancer activity. We found that most variants acted additively, while 30% showed interactions with other hSubs or with background variation, meaning that their effects were modulated by variants nearby. We also found pervasive additive and interactive background effects, indicating that segregating and chimpanzee-specific variants can have important consequences for enhancer activity differences between human and chimpanzee. The background effects we identified can obscure the effects of evolutionarily relevant hSubs, illustrating why it is important to distinguish the effects of segregating variants and fixed changes.

We found that differentially active fragments overall, and the effects of hSubs specifically, were not biased towards increased enhancer activity in human, but instead showed an even amount of bias towards either species. In principle, substitutions within HARs could increase or decrease enhancer activity. However, HGEs were defined using functional evidence of increased enhancer activity during human corticogenesis. Although HGEs were ascertained based on comparisons to rhesus macaque and mouse, we expect HGEs to include human enhancers that show increased activity relative to chimpanzee. Moreover, our studies focused on hSubs, which are derived sequence changes in human compared to an ancestral primate state shared between chimpanzee and rhesus macaque. In this context, we may not expect many HGE fragments to exhibit increased activity in chimpanzee compared to human. However, one potential explanation for this result is that our assays test individual fragments within HGEs that are components of much larger regulatory elements. In isolation, hSubs could increase or decrease activity, but in the context of the entire HGE may interact to increase activity overall. Our finding that hSubs and background variation interact in complex ways to alter activity at the fragment level provide support for this hypothesis.

We found enriched transcription factor binding site (TFBS) motifs among MPRA fragments active in our study, likely revealing transcription factors (TFs) that substantially contribute to gene regulation in human corticogenesis. However, we could not identify a subset of TFs associated with species-specific enhancer activity, suggesting that changes in regulatory function for the enhancers we studied are driven by the same set of TFs as those that drive enhancer activity in general. We did identify TFBSs that showed changes due to hSubs, and that were enriched in human versus chimpanzee active fragments. This supports that hSubs alter TFBS content in enhancers for specific transcription factors. Different TFBSs were uniquely enriched in human and chimpanzee fragments, implying that loss or gain of human enhancer activity due to hSubs involves changes in the recruitment of independent sets of TFs.

We also found that interacting hSubs were located significantly closer to each other than additive hSubs. This supports a model of TFBS evolution where variants interact by altering binding for the same TF or multiple TFs that bind in close proximity. For example, in the locus shown in Figures 6C–D, two close hSubs disrupt the predicted TFBSs of several transcriptional repressors, potentially underlying the higher activity of the human ortholog compared to chimpanzee. However, the small number of TFBS predictions overlapping multiple interacting hSubs precluded a systematic analysis.

Our study also suggests that HARs and HGEs encode regulatory changes with distinct evolutionary histories and potential biological effects. Notably, the effect sizes of hSubs in HGEs were on average larger than those in HARs. We had expected that hSubs in HARs might show larger effect sizes as they may be the result of positive selection for novel functions. Differences in the degree of sequence constraint between HARs and HGEs may explain our findings. HARs are highly constrained sequences, and remaining constraints on their ancestral functions may mitigate the effects of human-specific sequence changes within them. In contrast, HGEs are more weakly constrained, and many HGEs arose within placental mammals. Such young enhancers have been shown to have weakly conserved regulatory activity, and often exhibit changes in their activity across species (Berthelot et al., 2018; Cotney et al., 2013; Villar et al., 2015). The larger effect size we observed for hSubs in HGEs may reflect their increased evolutionary and functional plasticity compared to HARs. We also found that hSubs in constrained regions of HGEs had modestly larger effects than hSubs in unconstrained regions. We hypothesize that hSubs in constrained regions within HGEs may be altering ancestral regulatory functions, which may increase the activity of the pre-existing element. In contrast, hSubs in unconstrained regions may be giving rise to novel, but weak, enhancer activity, without a prior regulatory function to provide a “boost” for their effects.

Our finding that most hSub effects were modest supports that the evolution of uniquely human cortical features is in part the result of many genetic changes of small effect, consistent with the hypothesis that complex traits are highly polygenic (Boyle et al., 2017). Mammalian cortical development is a prime example of a polygenic trait affected by many loci of small effect, given the large numbers of genes involved which interact in complex gene regulatory networks (Geschwind and Rakic, 2013). However, it is important to note that the impact of an hSubs on enhancer activity in an MPRA may not reflect its impact in the native genomic context during cortical development. The hSubs we characterized here may have substantial biological effects despite their modest effects on enhancer activity in our MPRA. Further studies, including genetic models where the effects of hSubs can be determined *in vivo*, will be required to address this question.

We identified 532 HARs and HGEs with human-specific changes in enhancer activity in human neural stem cells, as well as individual sequence changes that contribute to those regulatory innovations. These findings now enable detailed experimental analyses of candidate loci underlying the evolution of the human cortex, including in humanized cellular models and humanized mice. Comprehensive studies of the HARs and HGEs we have uncovered here, both individually and in combination, will provide novel and fundamental insights into uniquely human features of the brain.

## Methods

### Target selection and initial MPRA library design

We selected genomic regions with potential human-specific regulatory activity by taking all Human Accelerated Conserved Noncoding Sequences (Prabhakar et al., 2006) and Human Accelerated Regions (version 2; Lindblad-Toh et al., 2011) (throughout the text collectively referred to as HARs) and all Human Gain Enhancers, that is, regions that show markedly higher ChIP-seq H3K27ac or H3K4me2 signal in human compared to mouse and rhesus macaque during corticogenesis (Reilly et al., 2015; Table S1). Within these regions, we collected all human–chimpanzee substitutions, that were fixed for the chimpanzee state in a primate alignment (comprising chimpanzee panTro4, orangutan ponAbe2, rhesus macaque rheMac3 and common marmoset calJac3) and likely to be fixed or nearly fixed in human populations (not present as a SNP marked as “Common” in dbSNP build 144, excluding indels). This resulted in a list of 32,776 human-specific Substitutions, or hSubs. We then designed 137 bp fragments centered on these hSubs. The fragment length was dictated by the maximal high-quality synthesis capacity at the time of 170 bp and the need to include primer sites for cloning. In case two hSubs were closely linked, we designed additional fragments centered on the midpoint between those hSubs. This way, we designed 49,629 experimental fragments per species. We included 639 control sequences per species from the same enhancer regions that did not contain any hSubs, for a total of 100,536 fragments. The final fragment library was synthesized by CustomArray.

## Experimental procedures for the first MPRA

Detailed descriptions of our MPRA library design, construction and experimental protocols are provided in the Supplemental Methods. Briefly, after synthesis, the fragment library was first amplified in a low-cycle PCR to complement the single synthesized strand. In a second step, fragments were tagged with random, 16 bp barcode tags in an emulsion PCR. The primer tails also contained restriction sites for cloning into the pMPRA1 vector and for cloning the *luc2* firefly luciferase ORF from pMPRAdonor2 in between fragment and barcode (Addgene plasmids #49349 and #49353 provided by T. Mikkelsen). After size selection using 2% agarose gels on a Pippin Prep (Sage Science #PL01717), the library was cloned into pMPRA1. This “inert” library was then sequenced on an Illumina HiSeq 2500 (2×250 bp) to establish fragment–barcode connections (see *Data analysis of the first MPRA*). We then cloned the luciferase ORF in between the fragment and the barcode to obtain the “competent” library used for the MPRA experiment.

### Cell culture and transfection

We used H9-derived human neural stem cells (hNSCs; GIBCO #10419428). These cells show a gene expression profile consistent with an early neurodevelopmental state, expressing *HES1*, *PAX6*, *NES* and *SOX2* according to RNA-seq (Cotney et al., 2015; see *Data and code availability*). They are multipotent and capable of differentiation into neurons, oligodendrocytes and astrocytes. hNSCs were grown in DMEM/F12 (GIBCO #12660012) with GlutaMAX (GIBCO #35050061), StemPro Neural Supplement (GIBCO #A1050801), bFGF (GIBCO #PHG0021) and EGF (GIBCO #PHG0311) added in flasks coated with Matrigel (Corning) at 37°C and 5% CO2. Cells were passaged using StemPro Accutase (GIBCO #A1110501) every 48 hours. We confirmed a uniform neural stem cell identity using immunofluorescent staining for SOX2 (Millipore #AB5603) and Nestin (Millipore #MAB5326) in parallel to transfection (Figure S7).

To perform the MPRA experiment, the library was transfected into hNSCs using a Nucleofector 2b (Lonza #AAB-1001). We used the plasmid’s luciferase activity to determine optimal experimental conditions (Supplemental Methods and Figure S8). Under final experimental conditions, we transfected four replicates (two each at hNSC passages 12 and 18) of five times 4 M cells with 32 µg of library per replicate and incubated cells for 6 hours before harvest with Accutase.

### Library harvest and sequencing

We used 95% of the harvested cells for total RNA extraction and the rest for plasmid DNA (pDNA) extraction. Initially, genomic DNA was extracted using the Qiagen AllPrep Kit, after which pDNA was enriched using the Qiagen MinElute PCR Purification Kit. RNA was purified using the Qiagen RNeasy Mini Kit including two DNase digests to minimize pDNA carryover. RNA integrity numbers were obtained using a 2100 Bioanalzyer (Agilent) and were high (> 8) in each replicate. We synthesized cDNA with the Invitrogen SuperScript III reverse transcription kit using a mixture between a custom, library-specific primer and oligo-dT primers (Table S9). qRT-PCR was performed to estimate the optimal number of PCR cycles to amplify the library for sequencing, so as to stop amplification in the exponential phase of PCR to avoid over-amplification (Figure S9). pDNA and cDNA libraries were then sequenced on an Illumina HiSeq 4000 (2×150 bp).

### Data analysis of the first MPRA

Inert library sequencing reads (containing fragment and barcode) were trimmed of low-quality bases using FastX Trimmer (options -Q 33 -f 10) and assembled into read pair contigs using PEAR (-q 33). Outer adapter sequences were removed using Cutadapt (-e 0.16). Finally, the inner adapter was removed with Cutadapt, leaving barcode tag and fragment sequence. Fragment sequences were mapped to the original library using Bowtie2 (--rg-id) to generate a map connecting barcode sequences with fragment identity. Experimental pDNA and cDNA sequencing reads (containing the barcode) were trimmed using FastX (-Q 33 -l 131 for read one, -Q 33 -l 126 for read two), oriented correctly (see scripts) and read pairs were assembled using PEAR (-q 33). Adapter sequences were removed using Cutadapt (-e 0.16). Resulting barcode sequences were summed up and annotated using the barcode–fragment map. Once summarized, data was analyzed using R. All of our scripts are publicly available (see *Data and code availability*).

After summarization, pDNA and cDNA counts were normalized by library size and log_2_ transformed. The resulting data is effectively normally distributed. Very small data values showed a Poisson-like distribution, which is why we excluded barcodes with an average pDNA barcode count across replicates below - 5.25 (Figure S2). We then normalized each cDNA value by its associated pDNA value to calculate each barcode’s “activity”. We summarized barcodes by fragment by calculating the median of all barcode values associated with a fragment in the pDNA fraction and the cDNA fraction, as well as the median of the cDNA/pDNA ratio (i.e., the fragment activity as defined above) for each fragment in each replicate. We also summarized replicates from the same cell batch further in two groups of replicates (“lineages”). Subsampling numbers of barcodes per fragment showed that correlations between replicates stabilized if a fragment had twelve or more barcodes (Figure S2), which we used as a cutoff for downstream analyses.

To test for fragment activity, we calculated the pDNA density distribution for each replicate (or group of replicates) and determined the point of maximal density (*µ*). This was found to be the most conservative distribution summary (see Figure S10) and also accounted for an apparent artifact in the activity distribution of one of the replicates. We then tested the distribution of each fragment’s cDNA barcodes against *µ* using a one-tailed *t* test to identify active fragments. We applied Benjamini-Hochberg (BH) multiple testing correction and accepted a fragment as active if it had a *P*_BH_ < 0.1 in at least two replicates or one lineage. We also applied a one-tailed Mann-Whitney *U* test to the two lineages to account for potential non-normal data distributions (Table S2). For the final set of reported active fragments, we further required that all fragments had a log_2_ activity > 0.1, averaged over all replicates, and that all replicates had a cDNA count larger than the pDNA count.

For differential activity testing, we included all orthologous fragment pairs that were measured in both species and active in at least one of the species. We tested for differential activity by applying two-tailed *t* tests of activity of the human allele versus that of the chimpanzee allele. A fragment pair was accepted as differentially active if it had a BH-corrected *P*_BH_ < 0.1 in at least two replicates or between the two grouped replicates. To account for non-normal data distributions, we also applied a Mann-Whitney *U* test to the two groups of replicates. The resulting fragments were selected for dissecting the effects of linked variants in MPRA round two. To report statistics for differentially active fragments, we further required that all replicates agreed in the direction of differential activity and that the average log_2_ fold change over all replicates was > 0.1.

### Design and experimental procedures for the second MPRA

Based on the results of the first MPRA, we designed a second MPRA library with two major components. First, we included all active fragments for replication (2,704 orthologous fragment pairs). Second, for all differentially active fragments, we designed artificial fragments that included all possible combinations of hSub states on both human and chimpanzee background sequences (14,963 fragments). In 972 loci (i.e., differentially active fragment pairs from the first MPRA), this library covered 1366 hSubs. After designing this library, we found 278 more “hSubs” that overlapped with “common” dbSNP (build 144) SNPs, which we excluded *post hoc*. All numbers given in this article exclude those SNPs. This library also contained additional positive and negative controls. Positive controls included fragments from enhancers that were previously reported to actively drive transcription in hNSCs (enhancers regulating *Nestin*, *SOX2* and *TLX*; Islam et al., 2015; Lothian and Lendahl, 1997; Zhou et al., 2014) and high confidence hNSC candidate enhancers overlapping phastCons elements (Siepel et al., 2005). Negative controls were generated by permuting enhancer sequences to random, inactive regions in the genome and then permuting phastCons elements within those enhancers to generate ‘pseudo-phastCons elements’ in ‘pseudo-enhancers’ on which MPRA fragments were designed. The library was synthesized by Agilent.

The experiment was performed as before with the following changes. We used 17 bp barcode tags. We synthesized cDNA using a custom, library-specific primer only, with comparable results to MPRA 1. We also included an additional no-RT control to ensure pDNA carryover contamination was at acceptable levels. Libraries were sequenced paired-end for 150 cycles on an Illumina NovaSeq 6000.

### Data analysis for the second MPRA

Mapping and summarization was done as in the first MPRA with the following changes. We mapped fragment sequences including their surrounding adapter sequences to force full-length mapping of highly similar sequences. We only accepted matches with >99% sequence identity and removed all barcode tags that were duplicated. We also increased stringency when testing for activity and differential activity relative to the initial screen. Fragments were defined as active at a BH-corrected *P* < 0.05 in a *t* test in both MPRA replicates. We tested the activity of experimental fragments against the distribution of negative controls (see above). This was necessary because the second library consisted solely of fragments with prior evidence of activity. While this would mean that more transcript is present in the sample, sequencing to a degree comparable between libraries leads to lower sequencing depth relative to the same fragments in the first library or to the pDNA library. This is in contrast to the first MPRA round, in which most fragments were expected to be inactive and their distribution could thus be used as a null distribution for identifying active fragments. For the same reason we did not require the cDNA value to be larger than the pDNA value. Testing for differential activity between species, we used two-tailed *t* tests between orthologous sequences and accepted a fragment as differentially active with *P*_BH_ < 0.05 in both replicates. All replicates of significant fragments agreed in the direction of species bias and showed an average log_2_ fold change > 0.1.

To identify hSub-specific enhancer effects, we applied an ANOVA test per locus (i.e., differentially active fragment from the first MPRA) with each hSub, the background variation, and their interactions as factors and the fragment activity as the response variable. Note that locus complexity, and thus model complexity, varied across fragments, from one hSub and no background variation up to seven hSubs with background. For each significant factor, we extracted its mean effect and its effect size in standard deviations according to Cohen 1988. For two-way interactions, we calculated effect size relative to the background state in which the factor showed its larger effect (i.e., of the two alterative states of the interacting hSub or background sequence). Significant hSubs were then annotated using comparative genomic data including hNSC H3K27ac ChIP-seq (Cotney et al., 2015; see *Data and code availability*), ATAC-seq (this study) and hNPC Hi-C (Rajarajan et al., 2018) data. ATAC-seq data was generated in triplicate according to Buenrostro et al., (2015) and sequenced on an Illumina HiSeq 2500 (2×100 bp). Reads were mapped using Bowtie2 (option -X 2000) and open chromatin regions were called using MACS2 (options -B --nomodel --shift -25 --extsize 50). Overlap with hNSC H3K27ac ChIP-seq is calculated using peak calls from Cotney et al. (2015).

### TFBS enrichment analysis

We downloaded all vertebrate transcription factor binding site (TFBS) models for transcription factors (TFs) expressed in hNSCs from the JASPAR core database (version 2014) and predicted binding sites in the human (GRCh37/hg19) and chimpanzee (panTro4) genomes using FIMO (option --max-stored-scores 10000000). We then overlapped the union of human and chimpanzee TFBS predictions with measured, active, differentially active, human-biased and chimpanzee-biased fragments and tested for enrichment in each set of elements using the next largest set as a background (i.e., enrichment within differentially active fragments compared to all active fragments, *et cetera*) using 10,000 resampling replicates. We also overlapped predicted human and chimpanzee TFBSs with all 1,183 hSubs in active MPRA fragments and tested for an enrichment of TFBSs within the human or chimpanzee TFBS predictions relative to the union of both. *P* values were BH-corrected.

### CRISPRi enhancer validation

We designed CRISPR guide oligos (Table S9) for three enhancers with species-specific activity that interact with genes expressed in hNSCs. Guide oligos were annealed and cloned into a plasmid containing dCas9-KRAB-T2a-GFP under the control of a UbC promoter (Thakore et al., 2015). hNSCs were transfected with plasmid using nucleofection, then sorted using an S3e Cell Sorter (Bio-Rad #1451006) after 96 hours to collect GFP-positive transfected cells. RNA was isolated using a Qiagen miRNeasy Mini kit and transcribed into cDNA using SuperScript III First-Strand Synthesis. Gene expression was quantified using LightCycler 480 SYBR Green I Master mix on a Roche 480 LightCycler II. The expression of each target gene was measured in triplicate 20 *µ*l reactions for each sample and normalized to the housekeeping gene *TBP*.

### Data and code availability

MPRA and ATAC-seq data have been deposited under GEO accession GSE140983. Additional data used in this study are deposited under GEO accession GSE57369 (hNSC RNA-seq and H3K27ac histone ChIP-seq).

The code used to analyze the data is deposited at GitHub: https://github.com/NoonanLab/Uebbing_Gockley_et_al_MPRA

## Supplemental Material

Supplemental Figures 1–10

Supplemental Tables 1–9

Supplemental Methods

## Acknowledgements

This work was supported by a grant from the National Institute of General Medical Sciences (GM094780) and funds from the Kavli Institute for Neuroscience at Yale University to J.P.N., a Research Fellowship (UE 194/1-1) from the Deutsche Forschungsgemeinschaft (DFG) to S.U., an NSF Graduate Research Fellowship (DGE-1122492) to A.A.K., and an Autism Speaks Dennis Weatherstone Predoctoral Fellowship to E.G. We thank M. Alderman III for proof-of-principle CRISPR knockout experiments and B. Lesch for access to the cell sorter.

## Contributions

J.G. and J.P.N. conceived of and designed the study with input from S.U.; J.G., S.U., A.A.K. and N.G. performed experiments; S.U. and J.G. analyzed the data; S.U., J.G. and J.P.N. interpreted the data; S.K.R., E.G., C.S. and J.C. provided materials; S.U. and J.P.N. wrote the manuscript with input from the other authors.

