## Supplemental Figures 1-10 for "Massively parallel discovery of human-specific substitutions that alter neurodevelopmental enhancer activity"

**Figure S1**. Cloning scheme to generate the MPRA library.
**Figure S2**. Correlations between replicates vs. barcode number.
**Figure S3**. Three examples of combinatorial loci.
**Figure S4**. Expanded version of Figure 4D showing all 444 fragments with significant hSub effects.
**Figure S5**. *P* value distribution for TFBS enrichment tests.
**Figure S6**. Extended Figure 6C showing all predicted TFBSs.
**Figure S7**. Nestin and SOX2 immunostaining confirms neural stem cell identity.
**Figure S8**. Optimization of transfection parameters based on the luciferase activity of the MPRA plasmid's *luc2* reporter gene.
**Figure S9**. qRT-PCR to estimate optimal cycle number for experimental barcode amplification for sequencing.
**Figure S10**. Comparison of three different summary statistics, Mean, Median and $\mu$.

**Figure S1**. Cloning scheme to generate the MPRA library. CRE indicates the MPRA fragment (Candidate Regulatory Element). The location of the barcode tag is shown as a green box. See Supplemental Methods for details.

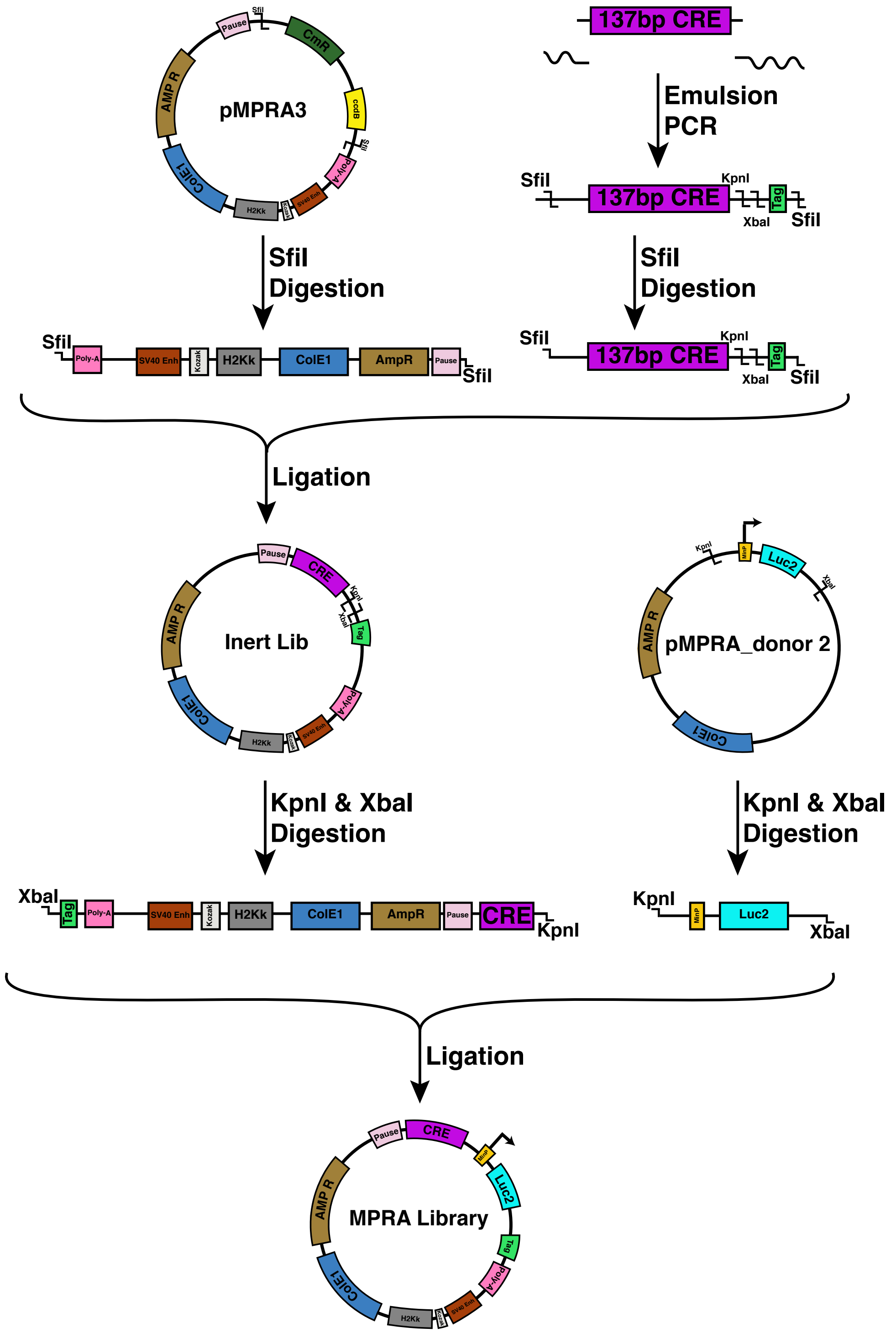

**Figure S2**. Correlations between replicates vs. barcode number. For all fragments having exactly N = 20, 40, 60, 80 or 100 barcode tags, two groups of barcodes per fragment were downsampled as N/2, (N/2) – 1, … , 2 barcode tags per group. Then the mean activity per fragment was calculated and Pearson's r was calculated between two sample replicates. By manual inspection of the inflection point at which the increase in correlation leveled off, we determined that fragments with twelve or more barcodes showed sufficiently high correlations to be included in downstream analyses.

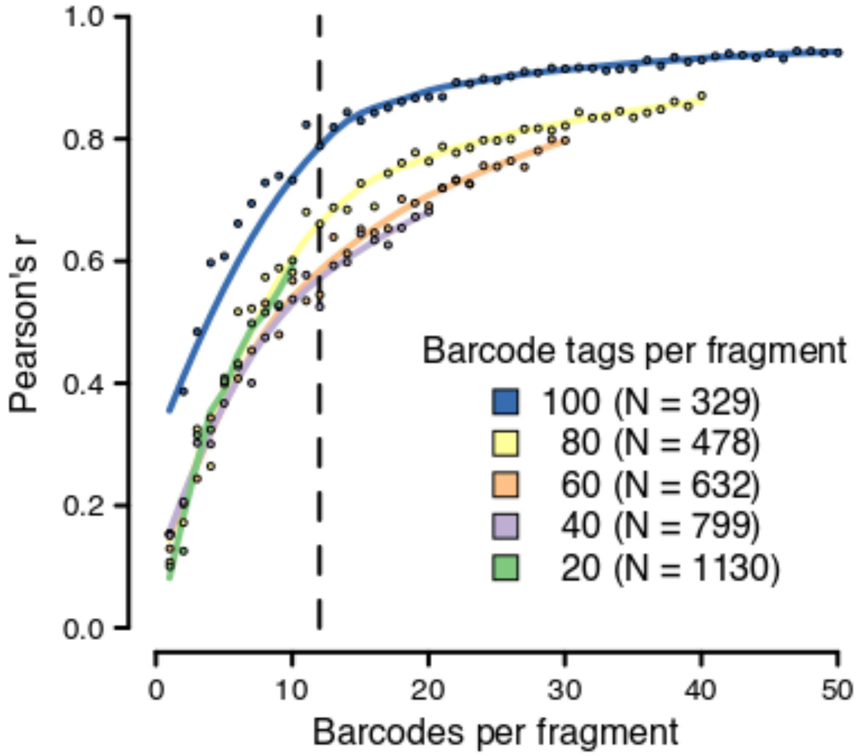

**Figure S3**. Three examples of combinatorial loci. Sketches (left) show the exact allele state. Filled blue boxes indicate the human hSub state, open green the chimpanzee state. Allele line color indicates background variation (blue, human; green, chimpanzee). Activity plots (right) show median and quartile ranges of fragment activity. (**A**) 2xHAR.63 contains the most human-biased locus with significant hSub effects in our dataset. The locus harbors four hSubs, two of which significantly affected regulatory activity in an additive manner. In addition, there was a slight but significant effect of background variation on activity. (**B**) HGE 2322 contains a locus that exemplifies visually additive hSub effects. This locus contains three hSubs, two of which showed significant effects on regulatory activity. Each of the two hSubs shown increases activity in the human allele state. Together in the same allele, these effects combine additively to produce the expected summed effect of both hSubs. There was no significant effect of the background variation. (**C**) HGE 2808 contains a locus that may serve as an example for an interactive effect. The locus contains three hSubs, two of which showed an effect on activity. Each of these two hSubs increases regulatory activity of the human allele slightly, but the combination of the two human states in the same allele increases activity more than expected by summing up their individual effects. The background variation present in this locus had no effect on activity.

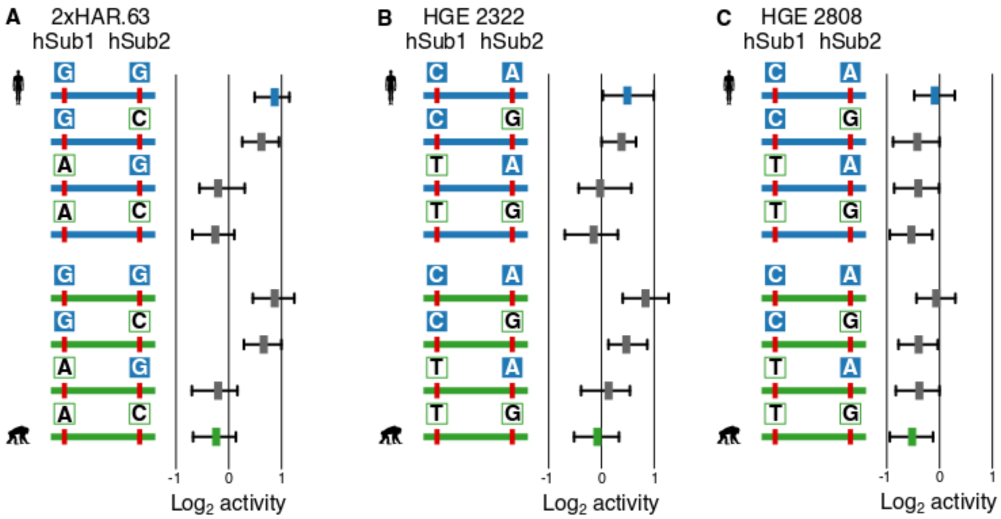

**Figure S4**. Expanded version of Figure 4D showing all 444 fragments with significant hSub effects. Fragments are sorted along the Y axis according to their fragment effect size from most human-biased to most chimpanzee-biased. Additive (blue) and interactive (orange) hSub effects are shown as dots in the same column.

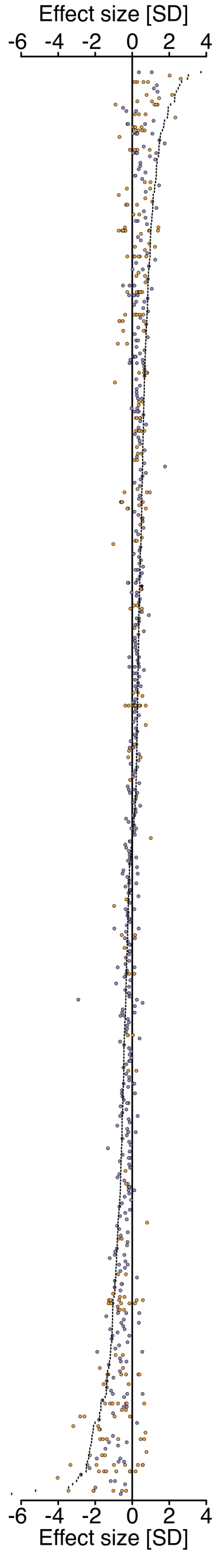

**Figure S5**. Uncorrected *P* value distributions for TFBS enrichment tests. See main text for details.

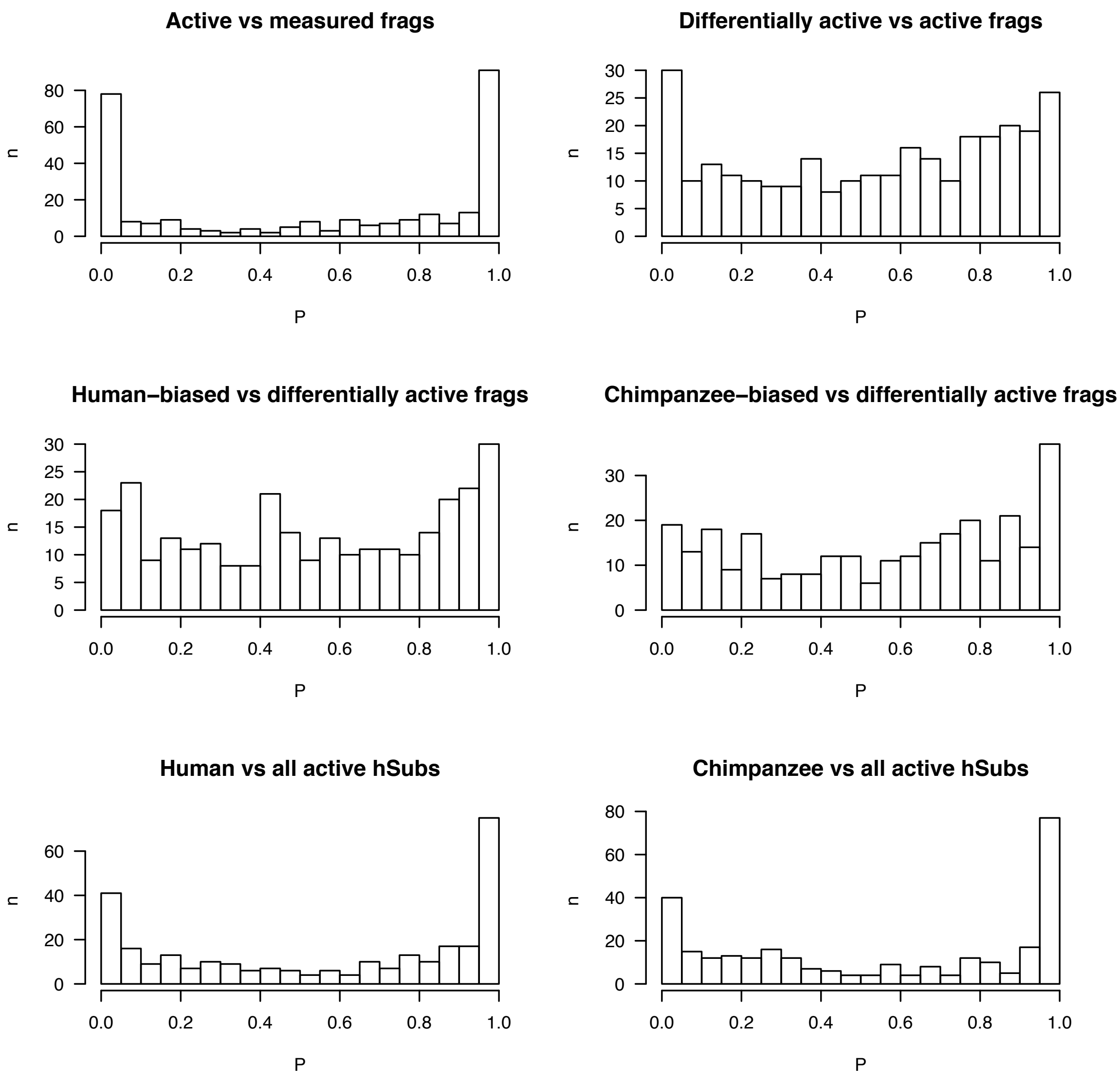

**Figure S6**. Extended Figure 6C showing all predicted TFBSs. Sites with stronger predicted binding in human than in chimpanzee are shown in blue, sites with stronger binding in chimpanzee are green, and sites predicted with similar binding strengths in both sequences are white.

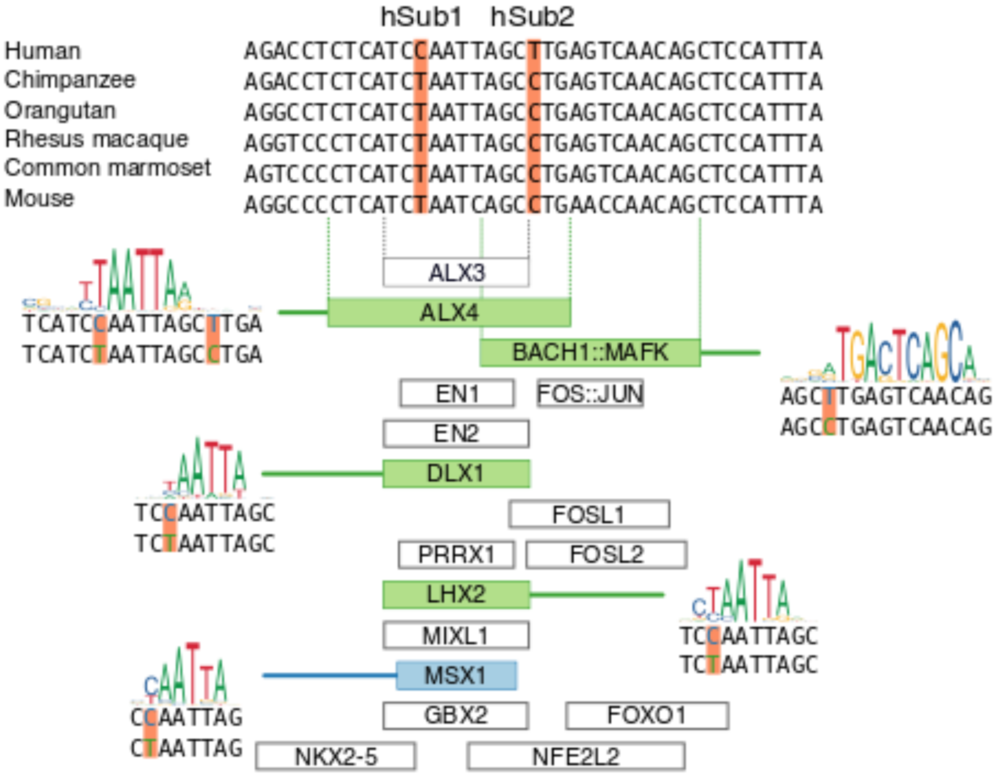

**Figure S7**. Nestin and SOX2 immunostaining supports neural stem cell identity. Staining was performed in parallel to each MPRA transfection.

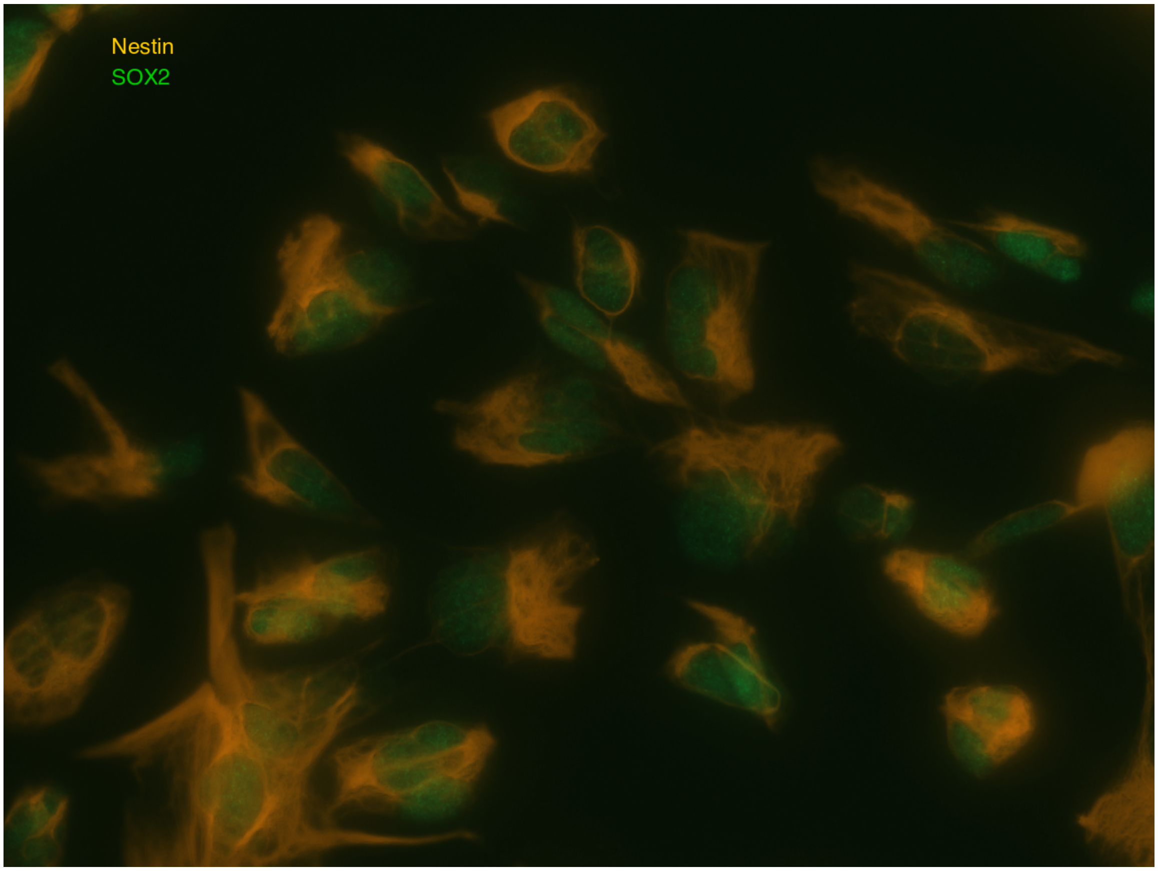

**Figure S8**. Optimization of transfection parameters based on the luciferase activity of the MPRA plasmid's *luc2* reporter gene. (**A**) Luciferase signal after varying incubation times. We had previously established an incubation time of < 12 hours as most effective. (**B**) qRT-PCR of the transcribed *luc2* gene to establish the optimal incubation time to harvest mRNA. This time is shorter than the time to maximal luciferase activity, potentially due to a translation time lag. (**C**) Transfecting varying amounts of plasmid per two million human neural stem cells. (**D**) Transfecting 16 $\mu$g of MPRA library with different numbers of cells to transfect. Note that the PGL3 control is extremely active in this experiment, while the negative control better reproduces previous results. Luciferase samples were standardized using 15 ng of co-transfected *Renilla* luciferase and luciferase signal is normalized relative to a vector containing the *PGL3* promoter. In **C–D**, cells were grown for 24 h after transfection.

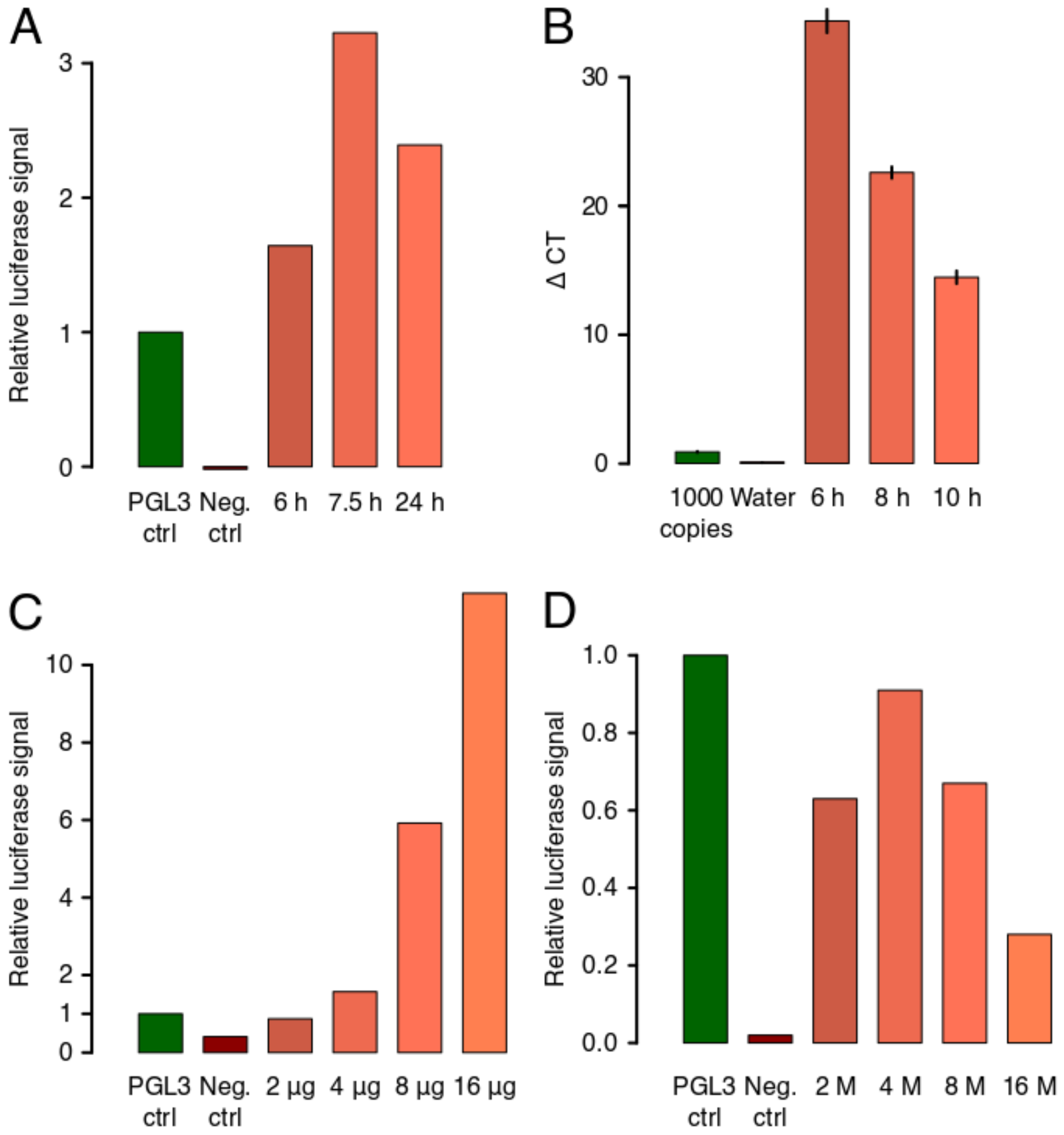

**Figure S9**. qRT-PCR to estimate optimal cycle number for experimental barcode amplification for sequencing. (**Top**) Amplification cycle number is first estimated in a full-length qRT-PCR run using a 10x dilution of pDNA and cDNA libraries. Given the target range, about 15 cycles should be sufficient for the undiluted library. (**Bottom**) qRT-PCR run of the undiluted library using 15 cycles shows amplification into the exponential phase. See Supplemental Methods for details

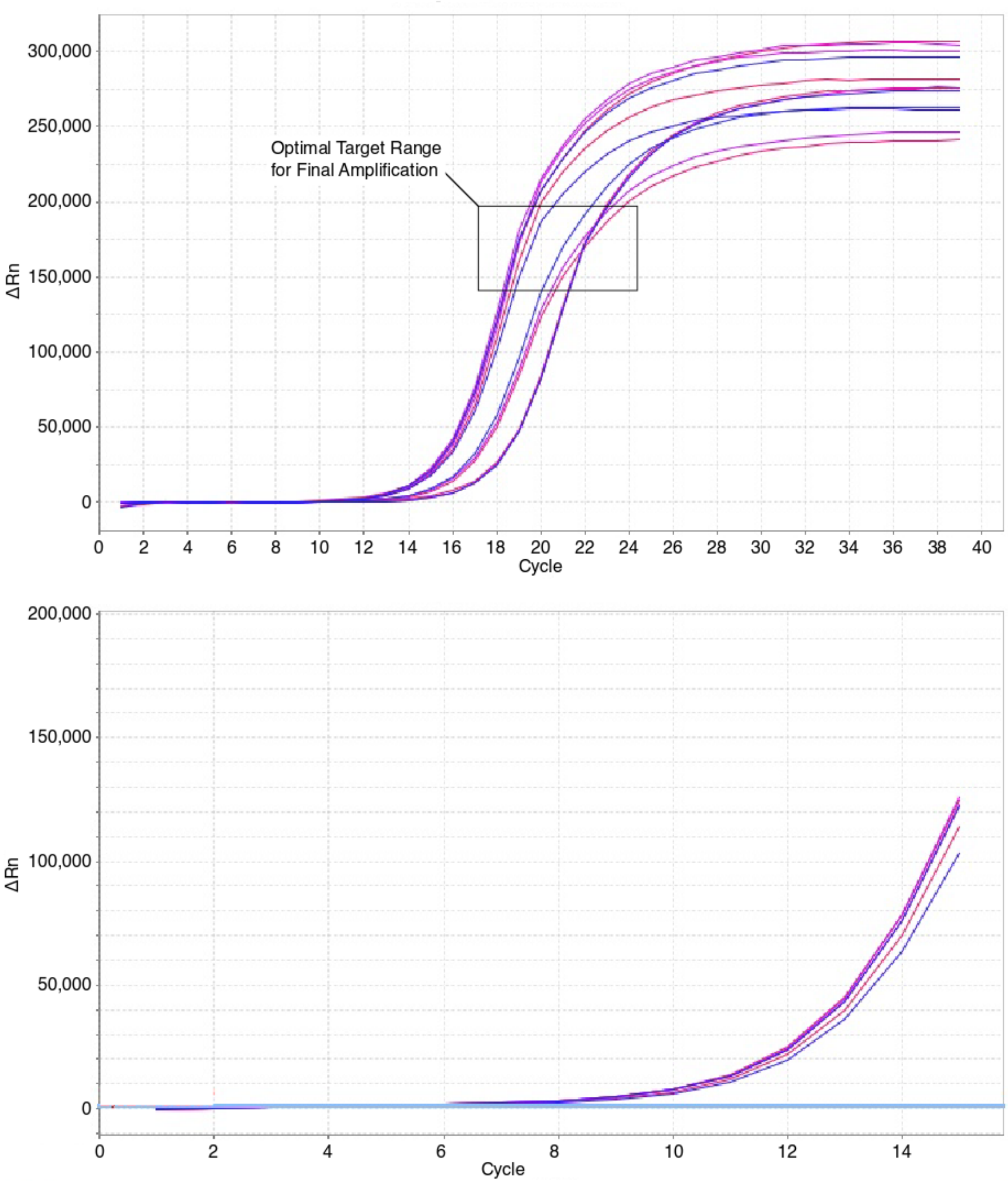

**Figure S10**. Comparison of three different summary statistics, Mean, Median and $\mu$. Numbers in the legend refer to unique fragments identified as active by the $t$ our line of tests dependent on the used summary statistic. (Note that the numbers reported in the main text are smaller due to additional constraints applied: Fragments had to have cDNA > pDNA in all replicates and an average activity >0.1.) We used $\mu$ as a measure of the distribution mean in our $t$ test in the first MPRA because it was the most conservative and because it accounts best for the ragged activity distribution of replicate 13.

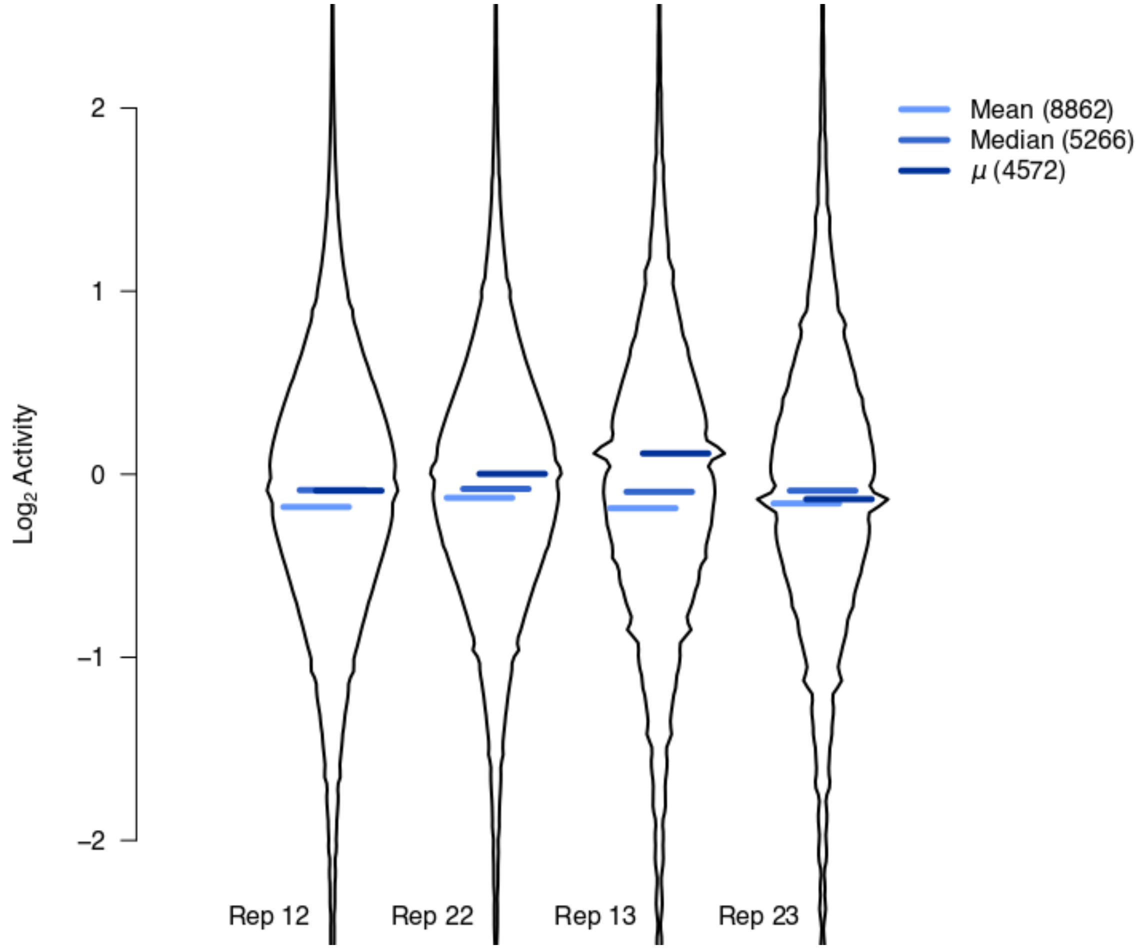
