## Supplemental Methods for "Massively parallel discovery of human-specific substitutions that alter neurodevelopmental enhancer activity"

### MPRA Supplemental Methods

November 19, 2019

### 1 Introduction

#### 1.1 Method Overview

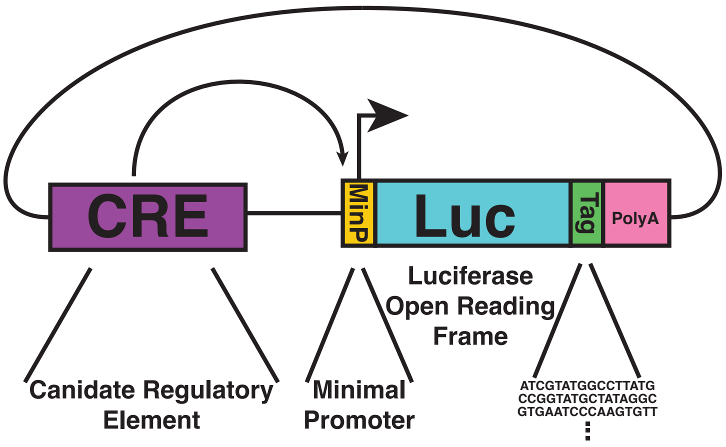

Figure 1: Mechanism of Enhancer Interrogation

.

- The MPRA assay relies on measuring candidate enhancers through their transcriptional capacities in parallel. A Candidate Response Element is cloned into a 5' orientation to a MinP driven ORF with a 16 random base pair tag between the ORF and poly-A tail. Utilizing the proportion of counts assigned to an individual transcript relative to that of the template DNA activity is measured as a fold enrichment.

#### 1.2 Library Construction Overview

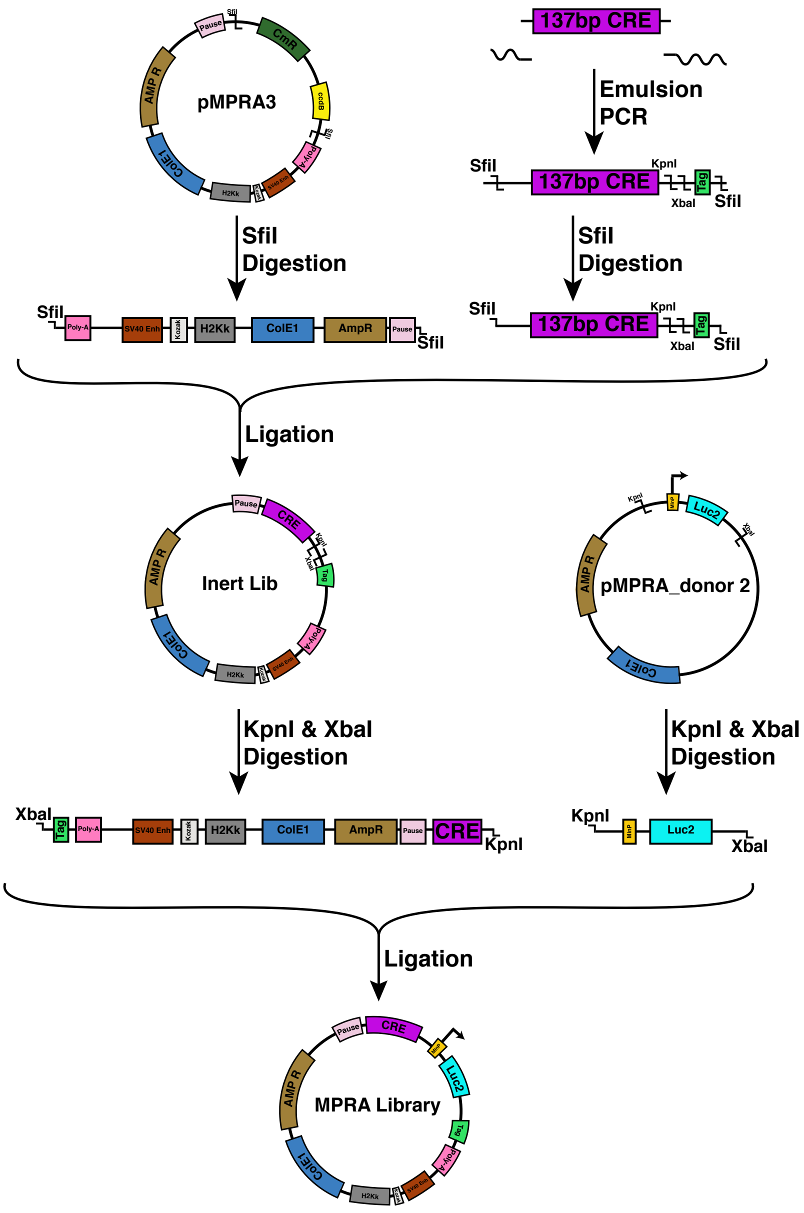

Figure 2: MPRA Library Construction

- The basic schematic of MPRA library construction above includes an emulsion PCR tailing reaction (table 7) and two cloning steps to create a competent library containing an active ORF driven by MinP.

##### 1.3 Emulsion PCR Overview

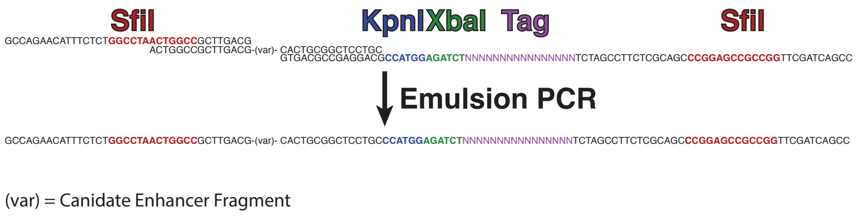

Figure 3: Tailing emulsion PCR

- The SfiI cloning arms are added to the distal ends to clone into the MPRA backbone vector, while the KpnI and XbaI sites lie in between the candidate enhancer and tag sequence. These sites are used to directionally clone in our Promoter-ORF complex.

### 2 Library Design

#### 2.1 Fragments

- Fragment size is limited by synthesis capacity. The maximum insert size is the maximum synthesis length minus 30 base pairs to account for incorporation of flanking amplification primers.

#### 2.2 Library Primers

- First set of primers are un-tailed
  Forward: 5' - GCCAGAACATTTCTCT - 3'
  Reverse: 5' - GCAGGAGCCGCAGTG - 3'

- Second set primers have tails used to add restriction sites and tags to synthesized fragments.

- The Forward primer adds SfiI (GGCCNNNNNGGCC) cloning site:

  5' - GCCAGAACATTTCTCT**GGCCTAACTGGCC**GCTTGACG - 3'

- The Reverse primer adds SfiI (GGCCNNNNNGGCC), KpnI(CCATGG), XbaI(AGATCT), and a 16bp tag sequence:

CCGACTAGCTT**GGCCGCCGAGGCC**CGACGCTCTTCCGATCTNNNNNNNNNNNNNNNN**TCTAGAGGTACC**GCAGGAGCCGCAGTG

- The reverse primer tag sequences are made by IDT and each **N** base pair is hand mixed with all nucleotides having an equal proportion of incorporation. The resulting Primer library is purified via HPLC (PAGE has low yield and could skew representation, standard desalting would allow truncated primer products to contaminate primer pool).

### 3 Library QC and Preliminary Amplification

The synthesized library needs to be PCR amplified to complement the synthesized single strand and to select only full length synthesis product for the emulsion PCR. The goal is to amplify the library with as few PCR cycles as possible to minimize amplification biases. To reach this goal, primer and library concentration in the PCR mix must be dialed in. Too much primer results in primer dimers, too few primer leads to no or incomplete amplification, too much library leads to over-saturation of the PCR. RT qPCR and Bioanalyzer analyses are used to estimate the needed concentrations.

#### 3.1 Single Reaction

- **Small Cycle qPCR Library Amplification**
- Observe library amplification with fluorescent SYBR Green qPCR reporting on every cycle

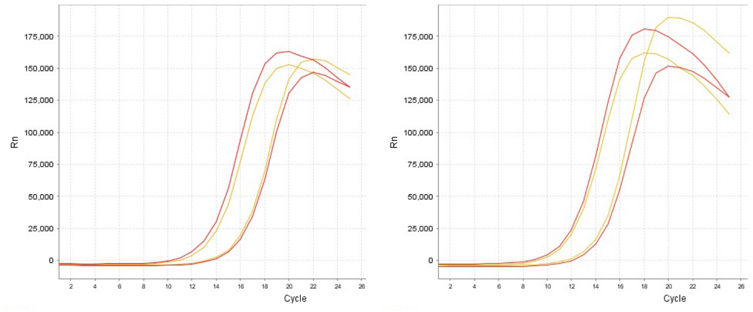

Figure 4: Initial qPCR Library Amplification Left: Human, Right: Chimp

- **PCR Mix:**

| Reagent | Volume | 30x |
|---|---|---|
| 2x NEB Master Mix | 10 µl | 300 µl |
| Water | 6.55 µl | 196.5 µl |
| Fwd + Rev primers (10 µmol/µl) | 1.25 µl | 37.5 µl |
| 10x SYBR Green | 1.2 µl | - |
| DNA 1:10 dilution | 1 µl | - |
| Final Volume | 20 µl | 600 µl |

Table 1: Initial library PCR.

- Pipette out 17.8 µl into 7 wells on qPCR plate.
- Add SYBR green and DNA to 5 wells (fluorescent reporter wells).
- Add SYBR green and water in place of DNA to 2 wells (control wells).
- Add water instead of SYBR green and and DNA to the 23 reaction master mix.
- Pipette 20 wells to collect for purification.
- **PCR Conditions:**

| Stage | Temperature | Time | Cycles |
|---|---|---|---|
| Stage 1 | 98°C | 30 Sec | 1 Cycle |
| | 98°C | 10 Sec. | |
| Stage 2 | 62°C | 30 Sec. | X Cycles |
| | 72°C | 30 Sec. | |
| Stage 3 | 72°C | 5 Min. | 1 Cycle |

Table 2: qPCR amplification conditions.

- Amplification cycles are determined empirically to stop amplification in the log phase.

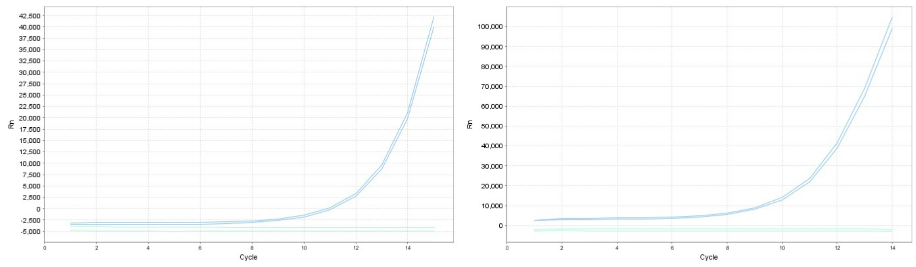

Figure 5: Left: Human, Right: Chimp

- Bead Purified 2x Volume ratio. Elute in 50 µl qiagen EB.

### 4 Emulsion PCR (CHIMERx: 3600-02)

#### 4.1 Reaction Mix

- **Emulsion:**

| Reagent | Volume | 16x |
|---|---|---|
| Emulsion Component 1 | 220 µl | 3520 µl |
| Emulsion Component 3 | 60 µl | 960 µl |
| Emulsion Component 2 | 20 µl | 320 µl |
| Final Volume | 300 µl | 4800 µl |

Table 3: Emulsion component mixture.

- Add components in order, use a wide bore tip for Emulsion Component 2.
- Incubate at 4°C for 30 minutes on wet ice.
- **Aqueous PCR Mix:**

| Reagent | Volume | 16x |
|---|---|---|
| 2x NEB Master Mix | 25 µl | 400 µl |
| Water | 19 µl | 304 µl |
| Fwd + Rev primers (5 µmol/µl) | 2.5 µl | 40 µl |
| BSA (10 mg/ml) | 2 µl | 32 µl |
| DNA | 1 µl | 16 µl |
| q5 Pol | 0.5 µl | 8 µl |
| Final Volume | 50 µl | 800 µl |

Table 4: Emulsion component mixture.

- Add entire volume to pre-chilled emulsion mix and vortex for 5 minutes on high at 4°C.
- **PCR Conditions:**

| Stage | Temperature | Time | Cycles |
|---|---|---|---|
| Stage 1 | 98°C | 30 sec. | 1 Cycle |
| | 98°C | 20 sec. | |
| Stage 2 | 72°C | 10 sec. | 15 Cycles |
| | 72°C | 15 sec. | |
| Stage 3 | 72°C | 5 min. | 1 Cycle |

Table 5: PCR conditions for library insert amplification.

- Each PCR reaction is 50 µl per well.
- **Cleanup:**
- Pool all reactions and add 1 ml of 2-butanol. Vortex thoroughly.
- Use kit provided spin columns.

#### 4.2 Full Scale Reaction

- **Emulsion:**

| Reagent | Volume |
|---|---|
| Emulsion Component 1 | 3520 µl |
| Emulsion Component 3 | 960 µl |
| Emulsion Component 2 | 320 µl |
| Final Volume | 4800 µl |

Table 6: Emulsion component mixture.

- Add components in order, use a wide bore tip for Emulsion Component 2.
- Incubate at 4°C for 30 minutes on wet ice.
- **Aqueous PCR Mix:**

| Reagent | Volume |
|---|---|
| 2x NEB Master Mix | 400 µl |
| Water | 319 µl |
| Fwd + Rev primers (5 µmol/µl) | 40 µl |
| BSA (10 mg/ml) | 32 µl |
| DNA | 1 µl |
| q5 Pol | 8 µl |
| Final Volume | 800 µl |

Table 7: Emulsion component mixture.

- Add entire volume to pre-chilled emulsion mix and vortex for 5 minutes on high at 4°C.
- **PCR Conditions:**

| Stage | Temperature | Time | Cycles |
|---|---|---|---|
| Stage 1 | 98°C | 30 sec. | 1 Cycle |
| | 98°C | 20 sec. | |
| Stage 2 | 72°C | 10 sec. | 15 Cycles |
| | 72°C | 15 sec. | |
| Stage 3 | 72°C | 5 min. | 1 Cycle |

Table 8: PCR conditions for library insert amplification.

- Each PCR reaction is 50 µl per well (96 wells total).
- **Cleanup:**
- Pool all reactions and add 13.7 ml of 2-butanol. Vortex thoroughly.
- Use kit provided spin columns. Condense all volume over 3–4 columns to concentrate eluted library.

#### 4.3 Size Select Library to Remove Slippage Products

- Slippage on the random tag sequence causes the production of additional products to arise from the emulsion PCR.

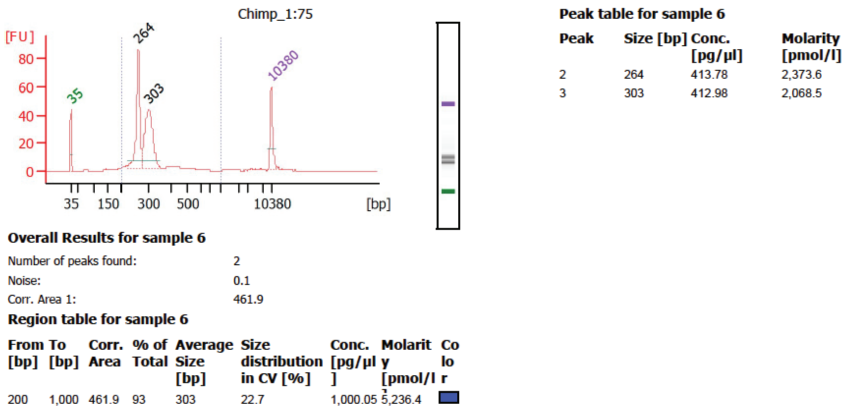

Figure 6: Extra large molecular weight slippage product ∼300 bp.

- Libraries were ran in a 2% Agarose pippen prep gel on Pippin Prep DNA Size Selection System to enrich for the correct 265 bp size product.

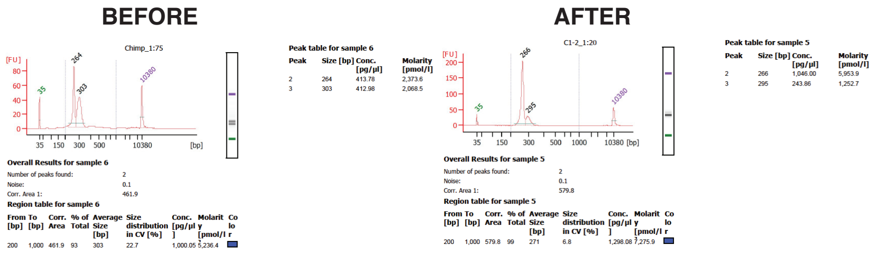

Figure 7: Tailed Libraries Before and After size selection.

### 5 Inert Library Cloning

#### 5.1 SfiI Digests

- Libraries and pMPRA1 vector were digested over night at 37°C. Digests were made as described in Table 9.

| Reagent | Library | pMPRA1 Vector |
|---|---|---|
| DNA, at least: | 200 ng | 700 ng |
| 10x Cutsmart buffer | X µl | X µl |
| SfiI | 2 µl/10 µl | 2 µl/10 µl |
| NcoI | - | 2 µl/10 µl |
| AclI | - | 2 µl/10 µl |
| XmaI | - | 2 µl/10 µl |
| Final Volume | X µl | X µl |

Table 9: Initial library digests.

- Add 2 µl CIP, 1 µl Cutsmart buffer and 7 µl water to the pMPRA1 vector digest. Incubate at 37°C for 1 hour.
- Clean up fragment library digests with MinElute columns. Elute in 8 µl EB buffer.
- The plasmid digest yields a 2.4 Kb band while all other bands are less than 600 bp. Use 0.5x AmpureXP beads to size select the DNA. This avoids a gel extraction.

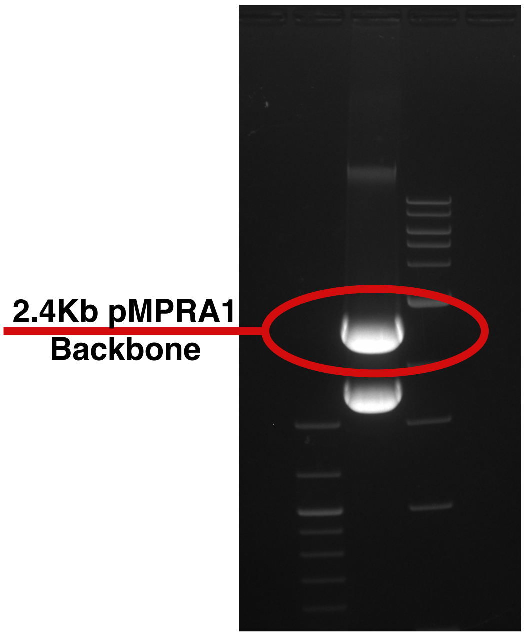

Figure 8: Gel extract pMPRA1 backbone to isolate from insert.

#### 5.2 Library Ligation

- Before setting up the ligation reaction confirm that the fragment libraries have been completely digested on both ends by running a Bioanalyzer high sensitivity chip on them.

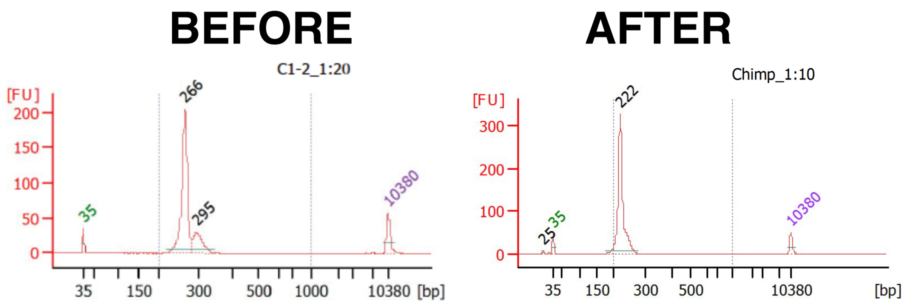

Figure 9: The 266 bp library fragments should lose ~40 bp after digest.

- Ligation reactions are performed at a fragment to backbone ratio of 3:1. Backbone input mass is 340 ng, making this particular library input mass 103 ng (corresponding to 3x backbone copy number). 2000 U of ligase is used per reaction. Ideally each reaction should yield at least 5,000,000 CFU. Reactions were setup as described in Table 10.

| Reagent | Amount |
|---|---|
| Water | ? µl |
| Fragment library | 103 ng |
| Vector backbone | 340 ng |
| 10x Ligation buffer | X µl |
| Ligase (400 U/µl) | 5 µl |
| Total Volume | X µl |

Table 10: Initial library ligation.

- Ligations incubated for 16 hours at 16°C in a thermocycler with 20–100 µl per tube.

#### 5.3 Library Transformation

- Ligations should be cleaned up with 1x room temperature AMPure XP beads. Add 1 volume of beads to the ligation reaction. Incubate for 5 minutes. Apply magnet for 5 minutes.
- Remove supernatant and wash twice with 200 µl 80% ethanol. Air dry for 10–15 minutes and elute in 10 µl EB.
- Label all LB-Amp plates (100 µg/ml) plates and place in 37°C incubator to pre-warm.
- Label tubes for a tenfold dilution series and fill with recovery media according to the serial dilution table below:

| Dilution | 1:10 | 1:100 | 1:1,000 | 1:5,000 | 1:10,000* | 1:50,000 | 1:100,000* |
|---|---|---|---|---|---|---|---|
| Recovery media | 90 µl | 90 µl | 90 µl | 40 µl | 90 µl | 40 µl | 90 µl |
| Cell volume | 10 µl | 10 µl | 10 µl | 10 µl | 10 µl | 10 µl | 10 µl |

Table 11: Serial dilution schema for ligation and transformation efficiency.
** These dilutions are made from the preceeding ten-fold dilution not the half fold dilution.*

- Place 5 electroporation cuvettes per 20 µl recombination reaction and a 1.5 ml Eppendorf tube on ice for each transformation reaction.
- Thaw cells on watery ice, make sure the water has time to come down to 4°C, mix cells by gently flicking the side of the tube. **DO NOT VORTEX!**
- Pipette 20 µl of thawed cells into the Eppendorf tube on ice with a **wide bore** pipette tip.
- Pipette 2 µl of DNA into the 20 µl of cells and tap tube to mix.
- Incubate on ice for 10 minutes.
- Take 21 µl of the cells with a **wide bore** pipette tip and place in the center of the cuvette. Tap the cuvette gently on the table to get the cells to fall into the bottom and rid the cells of any air bubbles.
- Place into electroporator arm and ensure that the metal cuvette is dry with a chem-wipe. Be sure not to warm the cuvette with your hand.
- Electroporate at 200 Ω, 25 µFd, 2.0 kV, and immediately place the cuvette back on ice.
- Place 1 ml of **room temperature** recovery into the cuvette.
- Use a **wide bore** 1 ml pipette tip to mix the cells once and then remove the cells slowly while tilting the cuvette on its side to ensure the cuvette well is drained. Place the cells into a 15 ml falcon round-bottom tube.
- Loosely fasten the cap to allow gas exchange and incubate at 37°C and 225 RPM for 1 hour.
- Pool all reactions and remove 10 µl. Dilute the cells according to Table 11. Plate two plates with 10 µl of each dilution.
- For every transformation, seed a **dense range** of 500 ml of LB-Amp in 2.5 l flasks. Allowing for a bit more than 20% variation, I chose 10, 15, 23, 36, 57, 90, 140, 220, 350, 550, 870, 1370, and 2150 µl.
- Grow inocculations at 37°C and 225 rpm for 5–11 hours and stop each inocculation at an $\mathbf{OD_{600} = 0.95\text{–}1.2}$. Check OD after 5 hours and from then on in regular intervals.
- First, remove 800 µl and transfer to a cryotube. Add 200 µl anhydrous glycerol, vortex to mix and store at -80°C to generate glycerol stocks.
- Pellet cells at 4500 rpm for 15 minutes at 4°C and freeze pellets at -20°C.

#### 5.4 Plasmid Library Purification

- Plasmid library is purified with HiSpeed Qiagen Maxi kit (12662).

#### 5.5 Estimate Complexity

- Calculate the concentration of competent transformants using the diluted series of plates equation:

$$CFUper\mu lseeded = \frac{AvgCFU \times DilutionFactor}{10\ \mu l}$$

- For a 52,000–104,000 element library, 80 tags per candidate enhancer is the target.
- Therefore for a 52,000 element library, total complexity is:

$$4,200,000 \approx CFUper\mu lseeded \times \mu lseeded$$

- Dilutions should be within 25% of your targeted complexity. It is better to err on the large end, because it is still possible to lose complexity afterwards. If you did not hit a satisfying complexity, repeat the transformation.

#### 5.6 QC Inert Library

- Equation to calculate total number of independent ligated transformants:

$$TotalComplexity = CFU \times DilutionFactor \times \frac{VolumeofCellsCultured(\mu l)}{10\mu l}$$

- Set up four different restriction digests of the purified inert library as described in table 12.

| Reagent | SfiI | XbaI | AfeI | NdeI |
|---|---|---|---|---|
| Water | 10 µl | 10 µl | 10 µl | 10 µl |
| DNA (100 ng/µl) | 5 µl | 5 µl | 5 µl | 5 µl |
| 10x Buffer | 2 µl | 2 µl | 2 µl | 2 µl |
| SfiI | 3 µl | - µl | - µl | - µl |
| XbaI | - µl | 3 µl | - µl | - µl |
| AfeI | - µl | - µl | 3 µl | - µl |
| NdeI | - µl | - µl | - µl | 3 µl |
| Total volume | 20 µl | 20 µl | 20 µl | 20 µl |

Table 12: Inert library digests.

- Run out all four digests on a 1% agarose gel.
- If library is correctly constructed it will digest according to Figure 5.6.

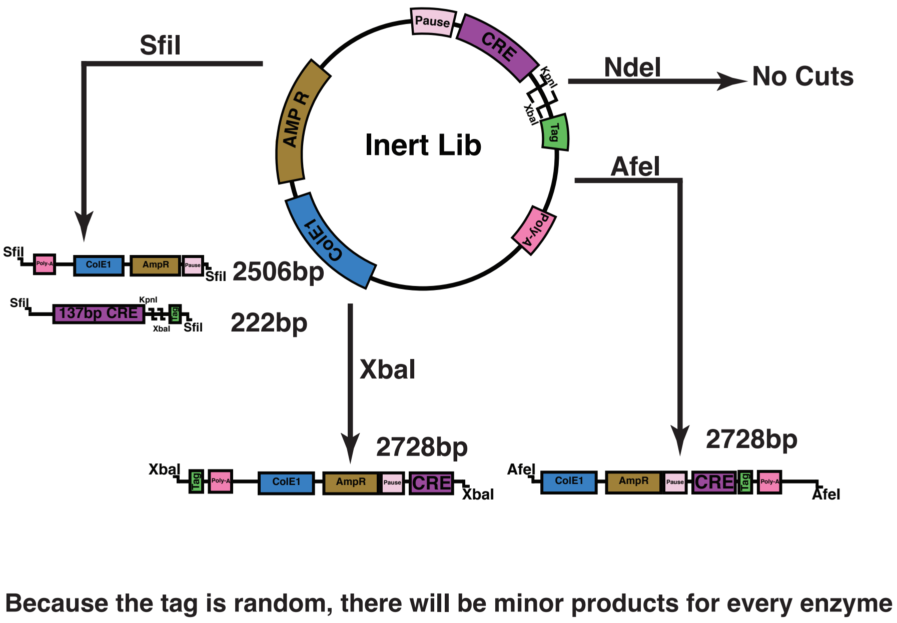

Figure 10: Library QC digest schematic

- Digestion gel should look similar to Figure 5.6.

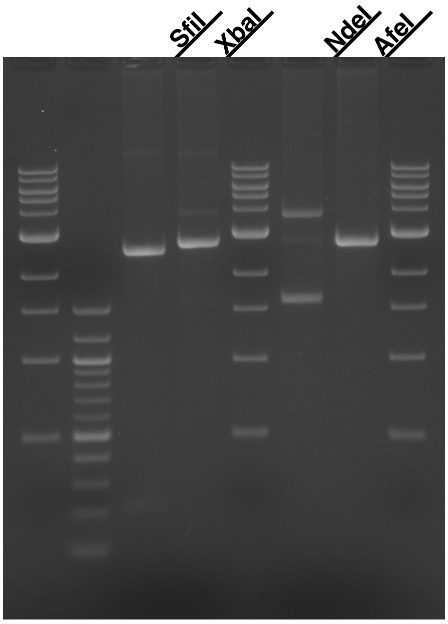

Figure 11: Library QC digest

### 6 Library Sequencing

#### 6.1 PCR Amplification of Inert Library

- Dilute primers to 100 µM, then mix forward and reverse primers to a final working concentration of 10 µM.
- Setup PCR reactions as follows:

| Reagent | 1x | 10x |
|---|---|---|
| 5X Q5 buffer | 10 µl | 100 µl |
| 5X GC enhancer | 10 µl | 100 µl |
| DMSO | 2.5 µl | 25 µl |
| Primers (10 µM) | 2.5 µl | 25 µl |
| dNTPs (25 mM) | 0.4 µl | 4 µl |
| Library DNA (3 ng/µl) | 0.5 µl | 5 µl |
| Q5 polymerase | 0.5 µl | 5 µl |
| Water | 22.25 µl | 236 µl |
| Final Volume | 50 µl | 500 µl |

Table 13: Volumes for library PCR reactions.

- Run 10 PCR reactions under the following conditions:

| Stage | Temperature | Time | Cycles |
|---|---|---|---|
| Stage 1 | 98°C | 30 sec. | 1 Cycle |
| | 98°C | 10 Sec. | |
| Stage 2 | 66°C | 20 Sec. | 18 Cycles |
| | 72°C | 20 Sec. | |
| Stage 3 | 72°C | 5 Min. | 1 Cycle |

Table 14: PCR conditions for library insert amplification.

- Pool the 10 PCR reactions and cleanup with 1.4x AMPureXP Beads or Qiagen PCR Purification Kit. Elute in 30 µl EB.
- Check size fraction and concentration on Bioanalyzer.

#### The following steps are done by the sequencing center:

#### 6.2 dA-Tailing

- Combine the following:

| Reagent | Volume |
|---|---|
| End-repaired DNA | 42 µl |
| 10x NEBNext end repair reaction buffer | 5 µl |
| Klenow Fragment (3'–5' exo-) | 3 µl |
| Final volume | 50 µl |

Table 15: dA tailing reaction.

- Incubate at 37°C for 30 minutes.
- Clean up with Qiagen QIAQuick PCR Purification kit. Elute in 25 µl EB

#### 6.3 Adaptor Ligation

- Combine the following:

| Reagent | Volume |
|---|---|
| dA-Tailed DNA | 25 µl |
| 5x NEBNext Quick Ligation Reaction Buffer | 10 µl |
| 15 µM NEBNext adaptors | 10 µl |
| Quick T4 DNA ligase | 5 µl |
| Final volume | 50 µl |

Table 16: Adaptor ligation reaction.

- Incubate at 20°C for 15 minutes.
- Clean up with Qiagen QIAQuick PCR Purification kit. Elute **twice** in 25 µl EB (final volume = 50 µl)

#### 6.4 Indexing PCR Amplification

- Dilute PCR primers to a final working concentration of 50 µM.
- Setup PCR reactions as follows:

| Reagent | 1x | 5x |
|---|---|---|
| 2x NEB High Fidelity | 12.5 µl | 62.5 µl |
| Library DNA | 1 µl | 5 µl |
| F-Primer (10 µM) | 1.25 µl | 6.25 µl |
| R-Primer (10 µM) | 1.25 µl | 6.25 µl |
| Water | 9 µl | 45 µl |
| Final vlume | 25 µl | 125 µl |

Table 17: Volumes library PCR reactions.

- Run 4 PCR reactions under the following conditions:

| Stage | Temperature | Time | Cycles |
|---|---|---|---|
| Stage 1 | 98°C | 30 Sec. | 1 Cycle |
| | 98°C | 10 Sec. | |
| Stage 2 | 65°C | 30 Sec. | 10 Cycles |
| | 72°C | 30 Sec. | |
| Stage 3 | 72°C | 5 Min. | 1 Cycle |

Table 18: PCR conditions for library insert amplification.

- Pool the 4 PCR reactions and cleanup with 0.9x AMPureXP beads. Elute in 20 µl EB.
- Check size and concentration on Bioanalyzer.
- Sequence 2x250 bp at high depth according estimated complexity in 5.6 dope in 5% PhiX to add complexity for cluster generation.

### 7 Cloning Promoter/ORF into Inert Vector

#### 7.1 KpnI and XbaI Library Digests

- Digest inert vector libraries with KpnI over night at 37°C as described in Table 19.

| Reagent | Volume |
|---|---|
| Plasmid library | 15 µl |
| Buffer | 2 µl |
| KpnI | 3 µl |
| Total volume | 20 µl |

Table 19: Inert library digest with KpnI.

- Cleanup the digestion with Qiagen Minelute column.
- Digest inert vector libraries with XbaI over night at 37°C as described in Table 20.

| Reagent | Volume |
|---|---|
| Plasmid library | 15 µl |
| Buffer | 2 µl |
| XbaI | 3 µl |
| Total volume | 20 µl |

Table 20: Inert library digest with XbaI.

- Add 1 µl CutSmart Buffer, 3 µl CIP, 4 µl water and incubate 45 minutes at 37°C.
- Cleanup the digestion with Qiagen Minelute column.

#### 7.2 KpnI and XbaI ORF Digests

- Digest donorMPRA 2 plasmid with KpnI and XbaI over night at 37°C as described in Table 21.

| Reagent | Volume |
|---|---|
| Plasmid library | 21 µl |
| Buffer | 3 µl |
| XbaI | 3 µl |
| KpnI | 3 µl |
| Total volume | 20 µl |

Table 21: DonorMPRA 2 digestion

- Run reaction on 0.8% agarose gel and gel purify the lower 1790 bp band.
- Cleanup the gel extraction with 1x AMPureXP beads.

##### 7.3 Library Ligation

- Ligate 500 ng of inert plasmid library to 650 ng of MinP/Luc2 ORF (2:1 Insert:Vector) as follows (Table 22):

| Reagent | Volume |
|---|---|
| Digested inert plasmid library | X µl = 500 ng |
| Digested MinP/Luc2 ORF | X µl = 650 ng |
| 10x ligase buffer | 2 µl |
| T4 DNA ligase (400 U/µl) | 5 µl |
| Water | X µl |
| Total volume | 20 µl |

Table 22: MPRA library ligation

- Incubate ligation reaction at 16°C for 16 hours in a thermocycler.
- Clean up ligation reaction with 1x AMPureXP beads and elute in 20 µl EB.

##### 7.4 Library Transformation

- Label all LB-Amp plates (100 µg/ml) plates and place in 37°C incubator to pre-warm.
- Label tubes for a tenfold dilution series and fill with recovery media according to the serial dilution table below:

| Dilution | 1:10 | 1:100 | 1:1,000 | 1:5,000 | 1:10,000* | 1:50,000 | 1:100,000* |
|---|---|---|---|---|---|---|---|
| Recovery Media | 90 µl | 90 µl | 90 µl | 40 µl | 90 µl | 40 µl | 90 µl |
| Cell Volume | 10 µl | 10 µl | 10 µl | 10 µl | 10 µl | 10 µl | 10 µL |

Table 23: Serial dilution schema for ligation and transformation efficiency.
***** *These dilutions are made from the proceeding ten-fold dilution not the half fold dilution.*

- Place 10 electroporation cuvettes per 20 µl recombination reaction and 10 1.5 ml Eppendorf tubes on ice for each transformation reaction.

- Thaw cells on watery ice, make sure the water has time to come down to 4°C, mix cells by gently flicking the side of the tube. **DON'T VORTEX!**
- Pipette 20 µl of thawed cells into the Eppendorf tube on ice with a **wide bore** pipette tip.
- Pipette 2 µl of DNA into the 20 µl of cells and tap tube to mix.
- Incubate on ice for 10 minutes.
- Take 21 µl of the cells with a **wide bore** pipette tips and place in the center of the cuvette. Tap the cuvette gently on the table to get the cells to fall into the bottom and rid the cells of any air bubbles.
- Place into electroporator arm and ensue that the metal cuvette is dry with a chem-wipe. Be sure not to warm the cuvette with your hand!
- Electroporate at 200 Ω, 25 µFd, 2.0 kV, and immediately place the cuvette back on ice.
- Place 1 ml of **room temperature** recovery into the cuvette.
- Use a **wide bore** 1 ml pipette tip to mix the cells once and then remove the cells slowly while rotating the cuvette on its side to ensure the cuvette well is drained. Place the cells into a 15 ml falcon tube.
- Loosely fasten the cap to allow gas exchange and incubate at 37°C and 225 RPM for 1 hour.
- Pool all reactions and remove 10 µl and dilute the cells according to table 23. Plate two plates with 10 µl of each dilution.
- For every 2 transformations, seed a 500 ml culture of LB-Amp in 2.5 l flasks (5 cultures total).
- Grow colonies at 37°C and 225 rpm for 8–11 hours **until OD = 1.5–2.0**.
- Pellet cells at 4500 rpm for 15 minutes at 4°C.

#### 7.5 Plasmid Library Purification

- Resuspend all pellets together and purify with Qiagen EndoFree Mega (or 4 replicates of a Maxi) Kit.
- Add one additional centrifugation after neutralization of pellet lysis; 17,900 x g for 10 minutes at 4°C.
- Elute in half the recommended amount of elution buffer.

#### 7.6 Estimate Transformation Efficiency

- Calculate total transformants utilizing the diluted series of plates equation:

$$CFUper\mu lseeded = AvgCFU \times DilutionFactor \times \frac{TotalVolumeofCells\mu l}{10\mu l}$$

#### 7.7 QC Competent Library

- Perform diagnostic digests as explained in Table 24. Incubate at 37°C overnight.

| Reagent | KpnI/XbaI | SexAI | SalI |
|---|---|---|---|
| Water | 8.5 µl | 11.5 µl | 11.5 µl |
| DNA(200 ngµl) | 3.5 µl | 3.5 µl | 3.5 µl |
| 10x Buffer | 2 µl | 2 µl | 2 µl |
| KpnI | 3 µl | - µl | - µl |
| XbaI | 3 µl | - µl | - µl |
| SexAI | - µl | 3 µl | - µl |
| SalI | - µl | - µl | 3 µl |
| Total volume | 20 µl | 20 µl | 20 µl |

Table 24: Competent library digests.

- The expected band sizes are shown in Figure 12.

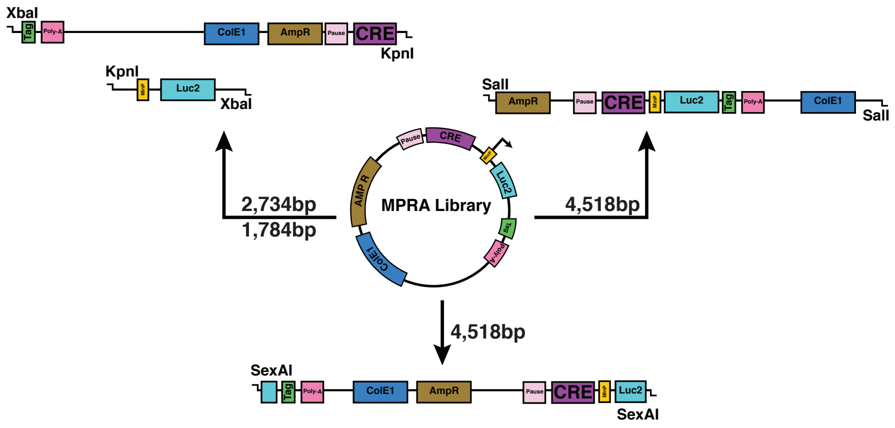

Figure 12: Competent library QC digest expected results

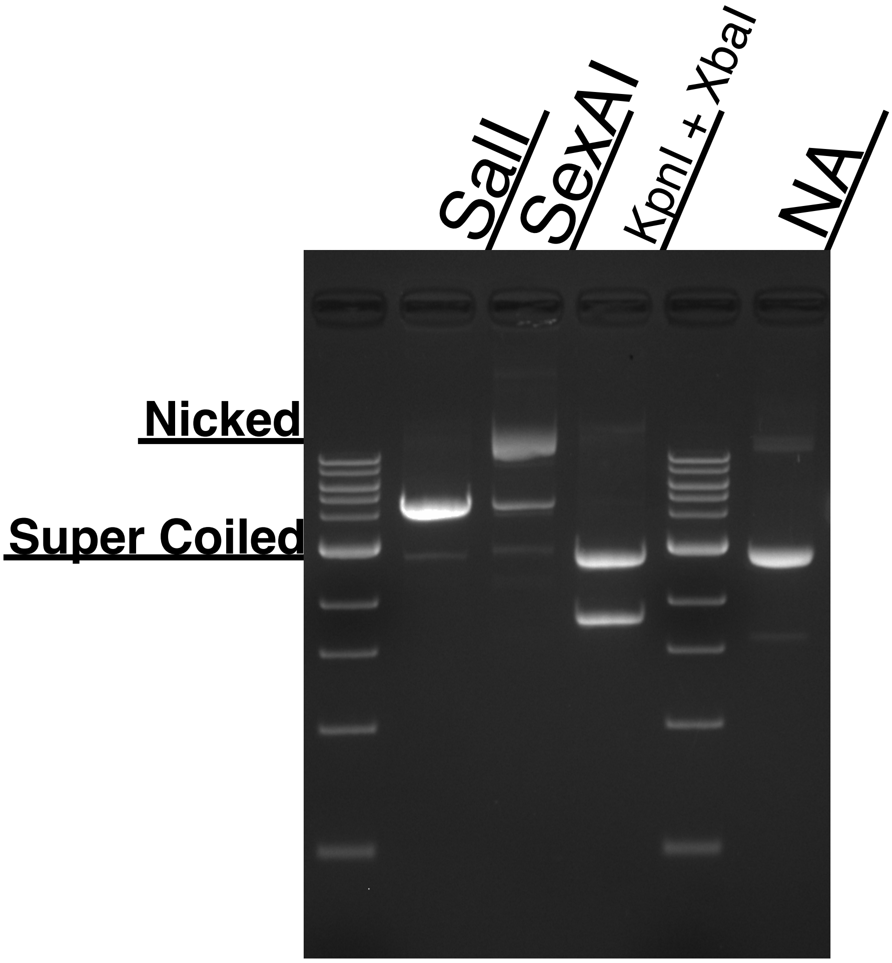

Figure 13: Competent library QC digest results

- Nicked and super-coiled variants may escape digestion therefore they can be identified by running non-digested plasmid in an alternate lane. In Figure 7.7, the super-coiled is around the same size as the backbone with MinP and luciferase removed (~2.7 kb) and SexAI appears to be low in activity.

#### 7.8 Barcode Sequencing of Competent Library

- PCR conditions for bar-code sequencing of competent library:

| Reagent | 1x | 9x |
|---|---|---|
| 2x NEB High Fidelity | 25 µl | 225 µl |
| Library DNA (15 ng/µl) | 1 µl | 9 µl |
| Fwd & Rev primer (10 µM) | 2.5 µl | 22.5 µl |
| Water | 21.5 µl | 193.5 µl |
| Final Volume | 50 µl | 450 µl |

Table 25: Volumes barcode PCR reactions.

| Stage | Temperature | Time | Cycles |
|---|---|---|---|
| Stage 1 | 98°C | 30 Sec. | 1 Cycle |
| Stage 2 | 98°C | 10 Sec. | 12 Cycles |
| | 65°C | 30 Sec. | |
| | 72°C | 30 Sec. | |
| Stage 3 | 72°C | 2 Min. | 1 Cycle |

Table 26: PCR Conditions for barcode amplification.

- Purify with Qiagen min elute and check for target amplicon size around 131 bp.
- Check size on bioanalyzer high sensitivity DNA chip.

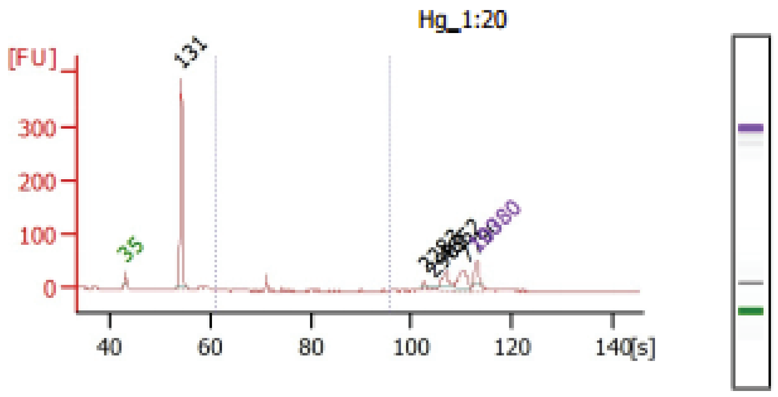

Figure 14: Bar-code containing amplicons

- Expected size with 5 degenerate bases on the 5' primer is 131 bp. Sequence amplicons 2x 150 bp.

### 8 Competent Library Characterization

#### 8.1 Library Activity

- Four co-transfections for each condition were performed at 2 µg of designated plasmid and 10 ng of PGL 4.72 Renilla luciferase per 2 million cells. 1 ml of media was added to each cuvette and 250 µl were plated per well on a six-well plate for 24 hours.
- Luciferase assays were performed with Promega Dual Luciferase Assays (E1910) and read on a Promega Luminometer (GM3500).

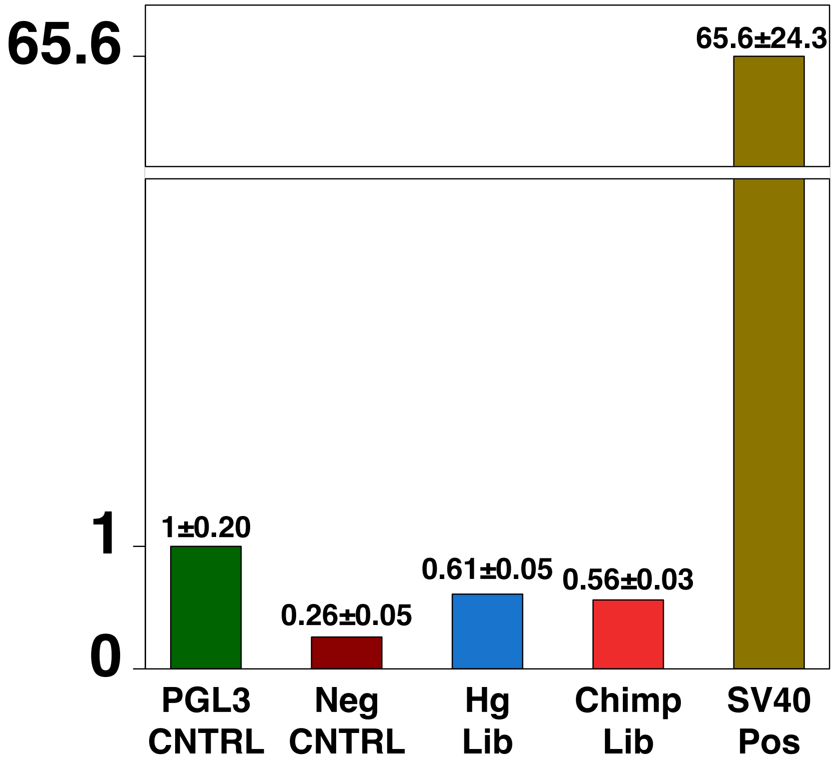

Figure 15: Competent Library Transfection

#### 8.2 Optimize Library Amount to Transfect

- Libraries were co-transfected with 2, 4, 8, and 16 µg of human plasmid library and 15 ng of 4.72 Renilla luciferase per 2 million cells. 1 ml of media was added to each cuvette and 250 µl were plated per well on a six-well plate for 24 hours.
- Luciferase assays were performed with Promega Dual Luciferase Assays (E1910) and read on a Promega Luminometer (GM3500).

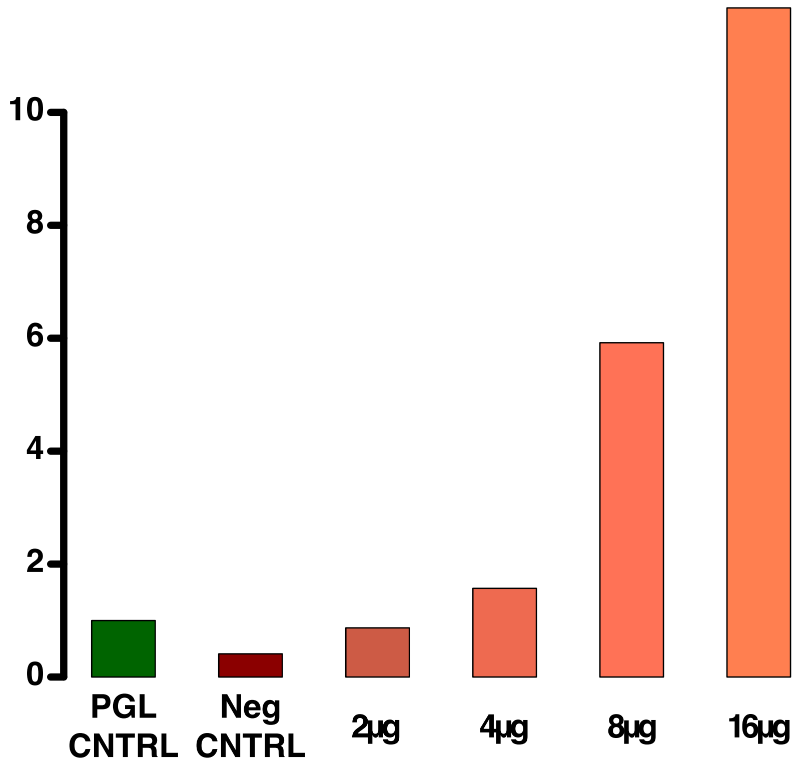

Figure 16: Transfection of increasing amounts of library.

#### 8.3 Optimize Amount of Cells to Transfect

- Libraries were co-transfected with 2, 4, 8, and 16 million cells, 16 µg plasmid library per 2 million cells, and 20 ng of 4.72 Renilla Luciferase per 2 million cells. 1 ml of media was added to each cuvette and 250 µl were plated per well on a six-well plate for 24 hours.

- Luciferase assays were performed with Promega Dual Luciferase Assays (E1910) and read on a Promega Luminometer (GM3500).

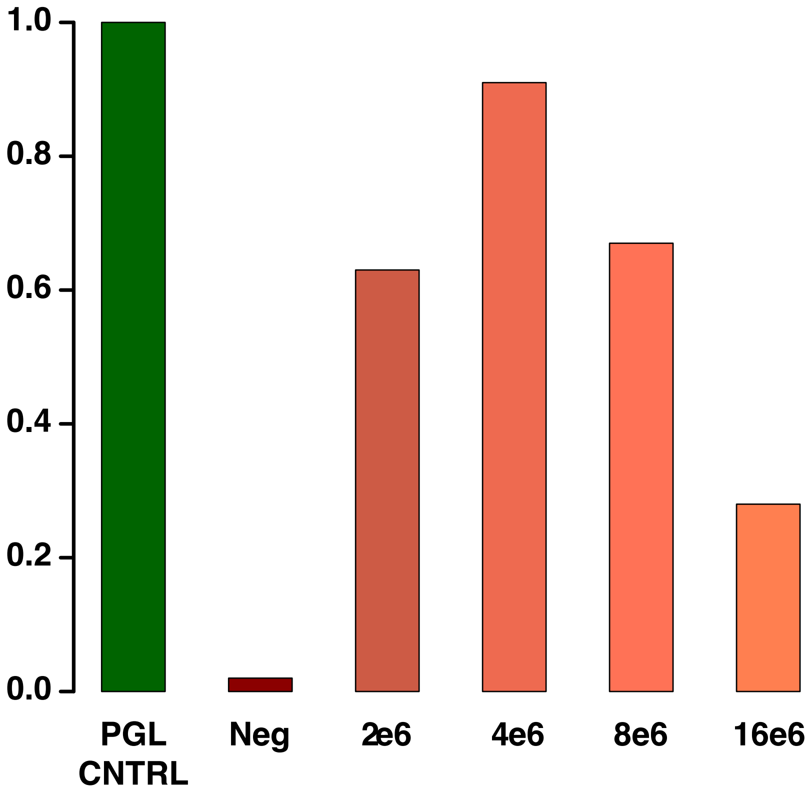

Figure 17: Transfection of Increasing Amounts Cells

- Note: PGL Control seems extremely active here. Around 10 fold more active than previously, as evidence by extremely low negative control activity.

### 9 Library Transfection

#### 9.1 Harvest the cells

- Wash a T150 flask of cells with 10 ml PBS(-), aspirate PBS.
- Pipette 8 ml pre-warmed Accutase into T150 flask and incubate for 5 minutes at 37°C.
- Tap the flask to dislodge cells and neutralize with 16 ml of complete media, pipette into a falcon tube and spin down cells at 200 x g for 4 minutes.
- Resuspend pellet in 5–10 ml of complete media.
- Count cells to quantify concentration.

##### 9.2 Transfection Conditions

- Pipette 2 x 20 million cells into 10 ml falcon tubes and spin down at 200 x g for 4 minutes.
- Aspirate media completely and pipette 2 x 160 µg of plasmid library on top of the cell pellet.
- Add 2 x 500 µl of Amaxa mNSC Reagent with Supplement 1 added and gently pipette to resuspend the cells.
- Gently resuspend the cell pellet and transfer to 2 x 5 Amaxa cuvettes.
- Electroporate the sample using pre-defined protocol **A-033**, and immediately add 1 ml of complete media to recover after each transfection.
- Use the pipette provided by Amaxa to transfer recovered cells into one falcon tube for both of the 5 x transfection rows.

##### 9.3 Plating Transfected Cells

- **Before transfection**: Coat two 150 $cm^2$ flasks each with 12 ml of coating media and one aliquot of matrigel for 1 hour to over night.
- Wash with 20 ml PBS(-), and add 30 ml of growth media. Leave plate in cell incubator to equilibrate.
- Add half of the pooled recovered cell volume to each equilibrated 150 $cm^2$ flask and grow for 6 hours before harvesting.

#### 10 Post Transfection Processing

##### 10.1 Harvest the cells

- Wash cells with 20 ml PBS(-).
- Pipette 10 ml pre-warmed Accutase into each 150 $cm^2$ flask and incubate for 6 minutes at 37°C.
- Tap the dish to dislodge cells and neutralize with 12 ml complete media, pipette into a 50 ml falcon tube and spin down cells at 200 x g for 4 minutes.
- Resuspend pellet in 10 ml PBS(-).
- Remove 5% of the total volume and place into a 1.5 ml Eppendorf tube for plasmid DNA library recovery. The remaining 95% are used for RNA extraction.
- Spin both the 10 ml falcon tube and the Eppendorf tube down at 200 x g for 4 minutes.

#### 10.2 Genomic DNA Purification

- Add 40 µl 1 M DTT per 1 ml of RLT buffer (Allprep kit).
- Aspirate PBS from the 1.5 ml Eppendorf tube and add 350 µl of previously prepared lysis buffer (RLT + DTT).
- Pipette to resuspend the cells and vortex vigorously to lyse cells.
- Spin the Eppendorf tube down and then add the entire volume to a Qiagen Allprep DNA/RNA Mini Kit column.
- Spin at 8150 x g for 30 seconds. **Proceed to Section 10.3 until instructed to return.**
- Wash column with 500 µl of buffer AW1 and spin at 8150 x g for 30 seconds.
- Wash column with 500 µl of buffer AW2 and spin at full speed for 2 minutes.
- Place in 1.5 ml Eppendorf tube, pipette 80 µl of EB onto the column, and incubate for 1 minute.
- Elute by spinning at 8150 x g for 1 minute.
- Pipette another 80 µl of EB onto the column and incubate for 1 minute.
- Elute a second time by spinning at 8150 x g for 1 minute.

#### 10.3 RNA Purification

- Add 40 µl 1 M DTT per 1 ml of RLT buffer and resuspend pellet in 600 µl of RLT lysis buffer plus DTT per 10 million cells transfected. Pipette to mix.
- Vortex well and pipette the solution onto a QiaShredder column in increments of 600 µl per column. Spin at max speed for 2 minutes.
- Add one volume of 70% EtOH to the flow through and pipette the combined solution onto an RNAeasy Column in increments of 650 µl. Spin at 8500 x g for 1 minute, this should equate to two spins per sample given this volume.
- Add 350 µl of RW1 buffer to the column and spin at 8500 x g for 30 seconds.
- Digest DNA on column by adding 10 µl pre-aliquoted DNAse I to 70 µl RDD buffer and pipetting all 80 µl directly onto the column. Incubate at room temperature with lid closed for 15 minutes. **While digesting the DNA return to Section 10.2 and finish the genomic DNA purification.**
- Add 350 µl of RW1 buffer to the column and spin at 8500 x g for 30 seconds.
- Add 500 µl of buffer RPE to the column and spin at for 30 seconds.
- Add 500 µl of buffer RPE to the column and spin at for 2 minutes.

- Place column in new collection tube and spin at full speed for 1 minute to dry.
- Place column in another new collection tube and pipette 30 µl of RNAse free water directly onto the column, wait 1 minute, and spin at 8500 x g.
- Pipette another 30 µl of RNAse free water directly onto the column, wait 1 minute, and spin at 8500 x g.
- Pool total RNA from the same transfections together and remove 2 µl aliquots of DNA for gel and nano-drop analysis.

#### 10.4 Second DNAse Treatment

- Prepare a second DNAse digestion as described in Table 27. Scale as necessary up to 200 µl, then split into different tubes.

| Reagent | 1x | 2x |
|---|---|---|
| RNA | Y µl | Y µl |
| Water | X µl | X µl |
| Buffer RDD | 10 µl | 20 µl |
| DNAseI | 2.5 µl | 5 µl |
| Final Volume | 100 µl | 200 µl |

Table 27: Second DNAse digestion mix.

- Incubate ad room temperature for 20 minutes.

#### 10.5 Plasmid Library Enrichment from Purified DNA

- **Perform while second digest is running in Section 10.4**.
- Add 300 µl ERC buffer to 20–100 µl of pDNA.
- Add 10 µl NaOAC (3M). Vortex to mix and spin down.
- Add total sample to QIAPrep Spin Miniprep column. Spin at 17,900 x g for 1 minute.
- Add 500 µl of PB buffer to column and spin at 17,900 x g for 1 minute.
- Add 750 µl of PE buffer to column and spin at 17,900 x g for 1 minute.
- Transfer column to new tube and spin at 17,900 x g for 1 minute to dry column.
- Place in 1.5 ml Eppendorf tube, add 30 µl EB onto the column, and incubate for 1 minute.
- Elute enriched pDNA by spinning at 17,900 x g for 1 minute.
- Pipette another 30 µl EB onto the column and incubate for 1 minute.
- Elute a second time by spinning at 8150 x g for 1 minute.

#### 10.6 RNA Clean-up

- Add 350 µl of RLT buffer for every 100 µl of DNAse-treated RNA.
- Add 250 µl of 96–100% RNAse free EtOH for every 100 µl of DNAse-treated RNA.
- Pipette up to 700 µl onto an RNeasy Mini Kit column and spin at 8500 x g for 30 seconds. Repeat as necessary up to column limit of 100 µg.
- Pipette 500 µl of RPE onto column and spin at 8500 x g for 30 seconds.
- Pipette 500 µl of RPE onto column and spin at 8500 x g for 2 minutes.
- Place column in new collection tube and spin at full speed for 1 minute to dry.
- Place column in another new collection tube and add 30 µl of RNAse free water directly onto the column. Wait 1 minute and spin at 8500 x g.
- Pipette another 30 µl of RNAse free water directly onto the column, wait 1 minute, and spin at 8500 x g.
- Pool total RNA from the same transfections and remove 2 µl aliquots of DNA for gel and nano-drop analysis.

#### 10.7 mRNA Purification using Dynabeads Kit (if Necessary)

- This step is usually not recommended due to unnecessary, high loss of RNA.
- Dilute 1–5 µg of total RNA to 50 µl with RNAse free water.
- Pipette 20 µl of resuspended beads in a **separate tube**.
- Add 100 µl of binding buffer to the beads and mix by pipetting gently.
- Place beads on magnet for 2 minutes.
- Remove supernatant, then remove tube from the rack and wash with another 100 µl of binding buffer. Mix gently by pipetting, place on magnet for 2 minutes, remove supernatant, remove tube from the rack.
- Resuspend the beads in 50 µl binding buffer and add 50 µl of total RNA. Mix gently by pipetting.
- Heat the entire reaction at 65°C for 5 minutes and then hold at 4°C or on ice for 3 minutes to denature secondary structure.
- Resuspend beads gently by pipetting.
- Incubate at room temperature for 5 minutes.
- Resuspend beads gently by pipetting.
- Incubate at room temperature for 5 minutes.

- Place tube on magnet for 2 minutes and discard supernatant.
- Remove tube from the rack and wash with 200 µl washing buffer. Mix by pipetting and place the tube on the magnet for 2 minutes. Discard supernatant.
- Repeat the previous step a second time.
- Resuspend beads in 50 µl of Tris, mix by pipetting, and transfer to thin wall 0.2 ml PCR tube.
- Heat sample in a thermocycler to 80°C for 2 minutes, then hold at 25°C to elute. Remove the samples when they reach 25°C.
- Add 50 µl binding buffer to each sample to allow the RNA to bind to the same beads. Mix by pipetting.
- Incubate at room temperature for 5 minutes.
- Resuspend beads gently by pipetting.
- Incubate at room temperature for 5 minutes.
- Place tube on magnet for 2 minutes and discard supernatant.
- Remove the tube from the rack and wash with 200 µl washing buffer, mix by pipetting then place the tube on the magnet for 2 minutes and discard supernatant.
- Spin down tube then place back on magnet and remove the remaining supernatant with a 20 µl pipette.
- Remove tube from the rack and resuspend beads in 17 µl Tris, mix by pipetting and transfer to thin wall 0.2 ml PCR tube.
- Heat sample in a thermocycler to 80°C for 2 minutes, then hold at 25°C to elute. Immediately remove samples when they reach 25°C and place on the magnetic rack for 2 minutes.
- Collect the enriched mRNA and remove 2 µl aliquots of DNA for gel and nano-drop analysis.

#### 10.8 RNA QC

- Run an Agilent Eukaryotic RNA Pico chip with samples of 1:100 and 1:1000 dilutions of extracted and digested RNA. If mRNA enrichment was performed, run lanes of mRNA and a 1:10 dilution of mRNA.

#### 10.9 cDNA Synthesis

- First strand synthesis performed with Invitrogen SSIII (18080-400). Reaction size scaled 2.5 x as follows.

| Reagent | Volume |
|---|---|
| Annealing buffer | 2.5 µl |
| Oligo dT | 2.5 µl |
| Custom primer | 2.5 µl |
| mRNA (no more than 5 µg) | 12.5 µl |
| Final volume | 20 µl |

Table 28: Volumes for a single oligo dT annealing reaction.

- Incubate 20 µl annealing reaction in DNAse/RNAse free PCR tube at 65°C for 5 minutes. Immediately place on ice.
- Add 2x buffer and SSIII enzyme to each reaction and incubate at 50°C for 2.5 hours in a thermocycler with the lid set to 60°C.

| Reagent | Volume |
|---|---|
| Annealing reaction | 20 µl |
| 2x buffer | 25 µl |
| SSIII enzyme | 5 µl |
| Final volume | 50 µl |

Table 29: Volumes for a single cDNA synthesis reaction.

##### 10.10 cDNA Second Strand Synthesis

- Add DEPC treated water, 10x buffer, and enzyme mix (NEB E6111) to each reaction. Pipette 50 µl aliquots into 0.2 ml PCR tubes and incubate at 16°C for 2.5 hours in a thermocycler with the lid left unheated.

| Reagent | Volume |
|---|---|
| 1st strand reaction | 50 µl |
| DEPC water | 120 µl |
| 10x buffer | 20 µl |
| Enzyme mix | 10 µl |
| Final volume | 200 µl |

Table 30: Volumes for a single cDNA synthesis reaction.

- Clean up reactions with Qiagen Minelute columns, try to keep volume as small as possible to help in PCR amplification.

#### 11 qPCR

##### 11.1 Preparation

- Primer stocks: All primers are kept at 100 µM, mix 10 µl forward with 10 µl reverse, and 80 µl for 100 µl of 10 µM working stock.
- Dilute 2 µl of cDNA 10-fold, and 100-fold.

- Dilute 2 µl of pDNA 10-fold, 100-fold, and 1,000-fold.
- Serial dilute plasmid standard out fresh each time from $1 \times 10^8$ copies / µl to $1 \times 10^2$ copies / µl.

#### 11.2 Master Mix

- All samples and standards are run in triplicate with an additional fourth reaction included to compensate for pipette error.
- Factor in 2–4 additional reactions into the master mix to account for pipette error as the master mix is very viscous.
- Include a water and genomic DNA control as well.
- Once total number of samples are accounted for make master mix as follows:

$$Samples_{total} = Samples \times 4 + 4$$

| Reagent | Volume/Rxn | Scale to volume |
|---|---|---|
| 2x Power Syber master mix | 10 µl | $\times Samples_{total}$ |
| DEPC water | 8 µl | $\times Samples_{total}$ |
| 10 µM F+R primer mix | 1 µl | $\times Samples_{total}$ |
| Sample | 1 µl | NA |
| Final volume | 20 µl | $19\mu l \times Samples_{total}$ |

Table 31: qPCR master mix.

- Pipette 76 µl into each well along the D or E rows of a 96-well Fast Optical qPCR Plate (Applied Biosystems 4346906).
- Add 4 µl of sample to each well.
- Use the multi-channel P200 pipette to add 20 µl of the 80 µl reaction into the three wells above (Row D) or below (Row E).

#### 11.3 Sample qPCR Plate Layout

- Below is a sample qPCR plate layout.

| | 1 | 2 | 3 | 4 | 5 | 6 | 7 | 8 | 9 | 10 | 11 | 12 |
|---|---|---|---|---|---|---|---|---|---|---|---|---|
| A | 1e8 | 1e7 | 1e6 | 1e5 | 1e4 | 1e3 | 1e2 | | | | | |
| B | 1e8 | 1e7 | 1e6 | 1e5 | 1e4 | 1e3 | 1e2 | | | | | |
| C | 1e8 | 1e7 | 1e6 | 1e5 | 1e4 | 1e3 | 1e2 | | | | | |
| D | Copy # Standard | | | | | | | | | | | |
| E | | | | | | | | pDNA | | cDNA | | Cntrl |
| F | | | | | | | | 1:100 | 1:1000 | 1:10 | 1:100 | GM | Water |
| G | | | | | | | | 1:100 | 1:1000 | 1:10 | 1:100 | GM | Water |
| H | | | | | | | | 1:100 | 1:1000 | 1:10 | 1:100 | GM | Water |

Master Mix

| | | | | | | |
|---|---|---|---|---|---|---|
| AOD 10uL/rxn | X | 56 | rxns. | = | 560 | uL |
| H2O 8uL/rxn | X | 56 | rxns. | = | 448 | uL |
| Prim 1uL/rxn | X | 56 | rxns. | = | 56 | uL |

Figure 18: Sample Layout of qPCR Plate

#### 11.4 Barcode PCR Amplification

- Generic PCR Master-Mix for Barcode-Seq:

| Reagent | Volume |
|---|---|
| 2x NEB Master Mix | 10 µl |
| Water | x µl |
| F+R primers (10 M) | 1.25 µl |
| DNA (20 million copies) | x µl |
| Final volume | 20 µl |

Table 32: Barcode PCR master mix.

- Generic PCR Conditions for Barcode-Seq:

| Stage | Temperature | Time | Cycles |
|---|---|---|---|
| Stage 1 | 98°C | 30 Sec. | 1 Cycle |
| | 98°C | 10 Sec. | |
| Stage 2 | 65°C | 20 Sec. | X Cycles |
| | 72°C | 30 Sec. | |
| Stage 3 | 72°C | 2 Min. | 1 Cycle |

Table 33: PCR conditions for sample barcode amplification.

- Before you can amplify the cDNA or pDNA fractions for sequencing you need to figure out the optimal amount of PCR cycles for however many template copies you want to seed for each 20 µl PCR reaction.
- Observation of amplification per-cycle (for 40 cycles) of the 1:10 dilutions with SYBR green under actual PCR conditions is needed as follows:

| Reagent | Volume |
|---|---|
| 2x NEB master mix | 10 µl |
| Water | 6.55 µl |
| F+R primers (10 µmol/µl) | 1.25 µl |
| 10x SYBR Green | 1.2 µl |
| DNA | 1 µl |
| Final volume | 20 µl |

Table 34: Barcode qPCR master mix.

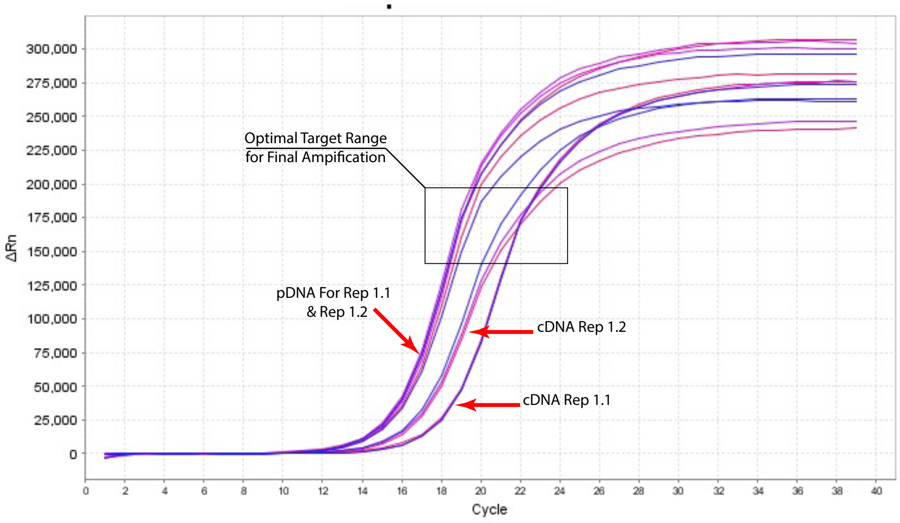

Figure 19: SYBR Green amplifiation

- Repeat amplification with undiluted pDNA. Take the cycle number around the desired range and subtract four cycles for the dilution change. Round up, and add one cycle to be safe. In the case of Figure 19 15 cycles seems sufficient.

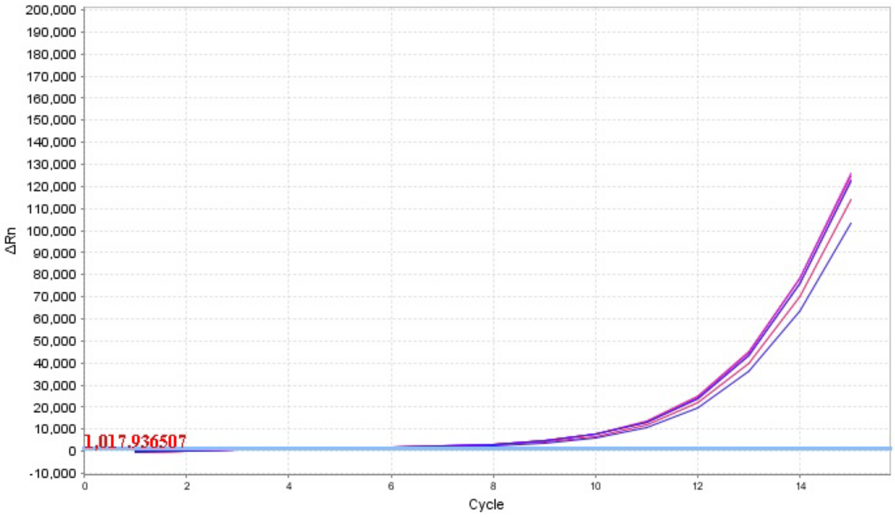

Figure 20: pDNA undiluted SYBR Green amplifiation

- Figure 20 is just on the lower end of optimal, but will work.
- Next step is to run out wells from both SYBR reactions on a 2% agarose gel and then purify seperate wells with a Qiagen Minelute column and run them on an Agilent DNA High Sensitivity chip. This will confirm that there are no off target effects and that there will be enough product produced at the flourescent intensity in Figure 20.
- Generic PCR Conditions for pDNA Barcode-Seq:

| Stage | Temperature | Time | Cycles |
|---|---|---|---|
| Stage 1 | 98°C | 30 Sec. | 1 Cycle |
| Stage 2 | 98°C<br>65°C<br>72°C | 10 Sec.<br>20 Sec.<br>30 Sec. | 17 Cycles |
| Stage 3 | 72°C | 2 Min. | 1 Cycle |

Table 35: PCR conditions for sample barcode amplification.

- Generic PCR Conditions for cDNA Barcode-Seq:

| Stage | Temperature | Time | Cycles |
|---|---|---|---|
| Stage 1 | 98°C | 30 Sec. | 1 Cycle |
| Stage 2 | 98°C<br>65°C<br>72°C | 10 Sec.<br>20 Sec.<br>30 Sec. | 18 Cycles |
| Stage 3 | 72°C | 2 Min. | 1 Cycle |

Table 36: PCR conditions for sample barcode amplification.

- PCR products are submitted to Illumina sequencing. Add 5% phi-X to help with sequence heterogeneity issues.

### 12 Reagents

#### 12.1 Consumables

| Reagent | Provider | Product number |
|---|---|---|
| Accutase, StemPro | Gibco | A11105-01 |
| Agarose | American Bio | AB00972-00500 |
| AMPureXP SPRI Beads | Beckman Coulter | A63880 |
| BSA (20 mg/ml) | NEB | B9000S |
| 2-Butanol | Acros Organics | 10770-0010 |
| Cells, MegaX DH10B T1$^{R}$ Electrocomp | Thermo Fischer Scientific | C6400-03 |
| Cuvettes for electroporation, 0.1 cm | Bio-Rad | 1652089 |
| DMSO | American Bio | AB03091-00100 |
| DNaseI, RNase-free set | Qiagen | 79254 |
| dNTPs | NEB | N0446S |
| Ethanol, 70% | American Bio | AB04010-00500 |
| Ethanol, 200 proof | American Bio | AB00515-00500 |
| Ethidium Bromide | Sigma | E1510-10ML |
| Glycerol, anhydrous | American Bio | AB00751-00500 |
| LB/Amp plates, 100 µg/ml | Recombinant Thechnologies | Stockroom |
| Ligase, T4 DNA (400 U/µl) | NEB | M0202M |
| Master Mix High Fidelity 2x | NEB | M0541L |
| NaOAC, 3M | American Bio | AB13168-01000 |
| PBS(-) (i.e., w/o MgCl & CaCl) | Gibco | 14190-144 |
| PGL 3 Control Vector | Promega | E174A |
| PGL 4.72 Renilla Luciferase Vector | Promega | E690A |
| Phosphatase, Alkaline (CIP) | NEB | M0290L |
| Pippin Prep gels, 2% agarose | Sage Science | CSD2010 |
| Q5 High fidelity polymerase | NEB | M0491L |
| qPCR Plates, fast optical | Applied Biosystems | 4346906 |
| SYBR Green | Life Technologies | S7563 |
| Water, DEPC | American Bio | AB02128-00500 |

Table 37: Primary consumable reagents.

#### 12.2 Kits

| Provider | Kit | Product number |
|---|---|---|
| Agilent | Bioanalyzer Eukaryotic Pico Kit | 5067-1513 |
| Agilent | Bioanalyzer High Sensitivity DNA | 5067-4626 |
| CHIMERx/EURx | Micellula Emulsion PCR Kit | 3600-02 |
| Invitrogen | Super Script III cDNA kit | 18080-400 |
| Lonza | Amaxa Mouse NSC Nucleofector Kit | VPG-1004 |
| NEB | NEBNext Library Kit (Seq center) | E6040L |
| NEB | HighSeq Indexing Primers (Seq center) | E7335L |
| Promega | Dual Luciferase Assay Kit | E1910 |
| Qiagen | AllPrep Mini DNA/RNA (for DNA) | 80204 |
| Qiagen | Endofree Mega Kit (before transfection) | 12381 |
| Qiagen | HiSpeed Maxi Kit | 12662 |
| Qiagen | Minelute | 28004 |
| Qiagen | QIAPrep Spin Miniprep Kit | 27106 |
| Qiagen | QIAshredder | 79654 |
| Qiagen | Reaction Clean-up (ERC Buffer is in here) | 28206 |
| Qiagen | RNeasy Mini Kit (for mRNA) | 74104 |
| Thermo Fisher | Dynabeads mRNA Purification Kit | 61006 |

Table 38: Primary kit reagents.

#### 12.3 Specialized Equipment

- Bio-Rad Gene Pulser
- Lonza Nucleofector 2b
- Applied Biosystems Step One Plus qPCR
- Agilent 2100 Bioanalyzer
- Promega Luminometer GM3500

### 13 Primer Sequences

#### 13.1 Initial Low Cycle Library Primers

- Forward: 5' - GCCAGAACATTTCTCT - 3'
- Reverse: 5' - GCAGGAGCCGCAGTG - 3'

#### 13.2 Library Bar-coding Primers

- Forward:5' GCCAGAACATTTCTCTGGCCTAACTGGCCGCTTGACG 3'
- Reverse:5' CCGACTAGCTTGGCCGCCGAGGCCCGACGCTCTTCCGATCTNNNNNNNNNNNNNNNTC-TAGAGGTACCGCAGGAGCCGCAGTG 3'

#### 13.3 Inert TagSeq Library Primers

- Forward:5' - (N:25252525)(N)(N)(N)(N)CAGGTGCCAGAACATTTCTCT - 3'
- Reverse:5' - TTATCATGTCTGCTCGAAGCGG - 3'

#### 13.4 Competent TagSeq Primer

- The 3' primer falls down of custom cDNA primer, allows for longer PCR product
- Forward:5' -(N:25252525)(N)(N)(N)(N)CAAGAAGGGCGGCAAGAT - 3'
- Reverse:5' - TTATCATGTCTGCTCGAAGCGG - 3'

#### 13.5 Barcode-Seq Primers (Targets cDNA)

- The 3' primer falls upstream of custom cDNA primer!!
- Forward:5' -(N:25252525)(N)(N)(N)(N)CGAGGTGCCTAAAGGACTG- 3'
- Reverse:5' - CCGACGCTCTTCCGATCT - 3'

#### 13.6 Reverse Transcription Primer

- 5' -CCGACTAGCTTGGCCGC- 3'

#### 13.7 Transcript qPCR Primers (targets Luciferase)

- Forward:5' -AACACCCCAACATCTTCGAC- 3'
- Reverse:5' -TCTCGGTCATGGTTTTACCG- 3'

#### 13.8 Backbone qPCR Primers

- Forward:5' -ATTTGGTATCTGCGCTCTGC- 3'
- Reverse:5' -TTTGCCGGATCAAGAGCTAC- 3'

#### 13.9 Experimental TagSeq Primer

- Forward:5' -(N:25252525)(N)(N)(N)(N)CGAGGTGCCTAAAGGACTG- 3'
- Reverse:5' -CCGACGCTCTTCCGATCT- 3'
